# A robust role for motor cortex

**DOI:** 10.1101/058917

**Authors:** Gonçalo Lopes, Joana Nogueira, George Dimitriadis, Jorge Aurelio Menendez, Joseph J. Paton, Adam R. Kampff

## Abstract

The role of motor cortex in non-primate mammals remains unclear. More than a century of stimulation, anatomical and electrophysiological studies has implicated neural activity in this region with all kinds of movement. However, following the removal of motor cortex, rats retain most of their adaptive behaviours, including previously learned skilled movements. Here we revisit these two conflicting views of motor cortex and present a new behaviour assay, challenging animals to respond to unexpected situations while navigating a dynamic obstacle course. Surprisingly, rats with motor cortical lesions show clear impairments facing an unexpected collapse of the obstacles, while showing no impairment with repeated trials in many motor and cognitive metrics of performance. We propose a new role for motor cortex: extending the robustness of sub-cortical movement systems, specifically to unexpected situations demanding rapid motor responses adapted to environmental context. The implications of this idea for current and future research are discussed.

## Introduction

The involvement of the brain and spinal cord in motor control has been recognized since the earliest known clinical records of head and spinal injuries, dating back to ancient Egypt (Louis, 1994; van Middendorp et al., 2010). However, the mechanism used by the nervous system to generate movement was not fully appreciated until Galvani first reported his famous experiments on *animal electricity* (Galvani, 1791). By isolating the sciatic nerve and gastrocnemius muscle in the frog, Galvani clearly demonstrated in a series of stimulation experiments that an electrical process, contained entirely within the biology of the frog’s leg, was responsible for the spontaneous generation of muscle contractions. This would lead to the discovery and physiological characterization of the nerve impulse, the action potential, that travels across the nerve to initiate muscle movement (du Bois-Reymond, 1843; Bernstein, 1868; Schuetze, 1983). The success of these seminal experiments immediately raised a fundamental question regarding nerve conduction: if spontaneous muscle contraction is generated by nerve impulses transmitted throughout the nervous system, how is this transmission coordinated in order to generate the complex patterns of muscle activity observed in natural behaviour?

### Discovery of the motor cortex

In search of answers to this question, researchers next turned to the brain, the seat of anatomical convergence of the nervous system. Following Galvani’s footsteps, several attempts were made to stimulate the cerebral cortex electrically, but with little success (Gross, 2007). It wasn’t until the 1870s that the first indications of a direct involvement of the cortex in the production of movement came to light, around the time when Hughlings Jackson undertook his studies on epileptic convulsions (Jackson, 1870). He observed that in some patients the fits would start by a deliberate spasm on one side of the body, and that different body parts would become systematically affected one after the other. He connected the orderly march of these spasms to the existence of localized lesions in the *post-mortem* brain of his patients and hypothesized that the origin of these fits was uncontrolled excitation caused by local changes in cortical grey matter (Jackson, 1870). In that same year, Fritsch and Hitzig published their famous study demonstrating that it is possible to elicit movements by direct stimulation of the cortex in dogs (Fritsch and Hitzig, 1870). Furthermore, stimulation of different parts of the cortex produced movement in different parts of the body (Fritsch and Hitzig, 1870). It appeared that the causal mechanism for epileptic convulsions predicted by Hughlings Jackson had been found, and with it a possible explanation for how the intact brain might control movement. The cerebral cortex was already considered at the time to be the seat of reasoning and sensation, so if activity over this so-called *motor cortex* was able to exert direct control over the musculature of the body, then it might, in the normal brain, be the area that connects volition to muscles (Fritsch and Hitzig, 1870).

### The Goltz-Ferrier debates

David Ferrier, a Scottish neurologist deeply impressed by the ideas of Hughlings Jackson and by the positive results of Fritsch and Hitzig’s experiments, proceeded to reproduce and expand on their observations with comprehensive stimulation studies showing how activity in the motor cortex was sufficient to produce a large variety of movements across a wide range of mammalian species (Ferrier, 1873). Meanwhile, other researchers across Europe such as Goltz and Christiani were facing a dilemma: in many of the so-called “lower mammals” massive lesions of the cerebral cortex failed to demonstrate any visible long-term impairments in the motor behaviour of animals (James, 1885; Goltz, 1888). These two lines of inquiry first clashed at the seventh International Medical Congress held in London in August 1881, where Goltz of Strassburg and Ferrier of London presented their results in a series of debates on the localization of function in the cerebral cortex (Phillips et al., 1984; Tyler and Malessa, 2000).

Goltz assumed a clear anti-localizationist position. He advanced that it was impossible to produce a complete paresis of any muscle, or complete dysfunction of any perception, by destruction of any part of the cerebral cortex, and that he found mostly deficits of general intelligence in his dogs (Tyler and Malessa, 2000). Following Goltz’s presentation, Ferrier emphasized the danger of generalizing from the dog to animals of other orders (e.g. man and monkey). He then proceeded to exhibit his own lesion results by means of antiseptic surgery in the monkey, describing how a circumscribed unilateral lesion of the motor cortex produced complete contralateral paralysis of the leg. He also produced a striking series of microscopic sections of Wallerian degeneration (Waller, 1850) of the “motor path” from the cortex to the contralateral spinal cord, the crossed descending projections forming the pyramidal corticospinal tract (Tyler and Malessa, 2000).

The debates concluded with the public demonstration of live specimens: a dog with large lesions to the parietal and posterior lobes from Goltz; and from Ferrier, a hemiplegic monkey with a unilateral lesion to the motor cortex of the contralateral side. As predicted, Goltz’s dog showed a clear ability to locomote and avoid obstacles and to make use of its other basic senses, while displaying peculiar deficits of intelligence such as failing to respond with fear to the cracking of a whip or ignoring tobacco smoke blown in its face. On the other hand, Ferrier’s monkey appeared severely hemiplegic, in a condition similar to human stroke patients. After the demonstrations, the animals were killed and their brains removed. Preliminary observations revealed that the lesions in Goltz’s dog were less extensive than expected, particularly on the left hemisphere. Ferrier’s lesions on the other hand were precisely circumscribed to the contralateral motor cortex. These demonstrations secured the triumph of Ferrier, who went on to firmly establish the localizationist approach to neurology and the idea of a somatotopic arrangement over the motor cortex.

The Goltz-Ferrier debates had far-reaching implications throughout the entire research community of the time, and the basic dilemma that was presented has sparked controversy and confusion for over a hundred years since (Phillips et al., 1984; Lashley, 1924; de Barenne, 1933; Tyler and Malessa, 2000; Gross, 2007). In the meantime, views of motor cortex have evolved to suggest it plays a role in “understanding” the movements of others (Rizzolatti and Craighero, 2004), imagining one’s own movements (Porro et al., 1996), or in learning new movements (Kawai et al., 2015), but where are we today regarding its role in directly controlling movement?

### Stimulating motor cortex causes movement; motor cortex is active during movement

Motor cortex is still broadly defined as the region of the cerebral hemispheres from which movements can be evoked by low-current stimulation, following Fritsch and Hitzig’s original experiments in 1870 (Fritsch and Hitzig, 1870). Stimulating different parts of the motor cortex elicits movement in different parts of the body, and systematic stimulation surveys have revealed a topographical representation of the entire skeletal musculature across the cortical surface (Leyton and Sherrington, 1917; Penfield and Boldrey, 1937; Neafsey et al., 1986). Electrophysiological recordings in motor cortex have routinely found correlations between neural activity and many different movement parameters, such as muscle force (Evarts, 1968), movement direction (Georgopoulos et al., 1986), speed (Schwartz, 1993), or even anisotropic limb mechanics (Scott et al., 2001) at the level of both single neurons (Evarts, 1968; Churchland and Shenoy, 2007) and populations (Georgopoulos et al., 1986; Churchland et al., 2012). Determining what exactly this activity in motor cortex controls (Todorov, 2000) has been further complicated by studies using long stimulation durations in which continuous stimulation at a single location in motor cortex evokes complex, multi-muscle movements (Graziano et al., 2002; Aflalo and Graziano, 2006). However, as a whole, these observations all support the long standing view that activity in motor cortex is involved in the direct control of movement.

### Motor cortex lesions produce different deficits in different species

What types of movement require motor cortex? In humans, a motor cortical lesion is devastating. Permanent injury to the frontal lobes of the brain by stroke or mechanical means is often followed by weakness or paralysis of the limbs in the side of the body opposite to the lesion (Louis, 1994). Although the paretic symptoms have a tendency to recover partially, especially with training and rehabilitation, permanent movement deficits and loss of muscle control in the affected limbs is the common prognosis; movement is permanently and obviously impaired (Laplane et al., 1977; Kwakkel et al., 2003). In non-human primates, similar gross movement deficits are observed after lesions, albeit transiently (Leyton and Sherrington, 1917; Travis, 1955). The longest lasting effect of a motor cortical lesion is the decreased motility of distal forelimbs, especially the control of individual finger movements required for precision skills (Leyton and Sherrington, 1917; Darling et al., 2011). But equally impressive is the extent to which other movements fully recover, including the ability to sit, stand, walk, climb and even reach to grasp, as long as precise finger movements are not required (Leyton and Sherrington, 1917; Darling et al., 2011; Zaaimi et al., 2012). In non-primate mammals, the *absence* of lasting deficits following motor cortical lesion is even more striking. Careful studies of skilled reaching in rats have revealed an impairment in paw grasping behaviours (Whishaw et al., 1991; Alaverdashvili and Whishaw, 2008), comparable to the long lasting deficits seen in primates, but this is a limited impairment when compared to the range of movements that are preserved (Whishaw et al., 1991; Kawai et al., 2015). In fact, even after complete decortication, rats, cats and dogs retain a shocking amount of their movement repertoire (Goltz, 1888; Bjursten et al., 1976; Terry et al., 1989). If we are to accept the simple hypothesis that motor cortex is the structure responsible for “voluntary movement production”, then why is there such a blatant difference in the severity of deficits caused by motor cortical lesions in humans versus other mammals? With over a century of stimulation and electrophysiology studies clearly suggesting that motor cortex is involved in many types of movement, in all mammalian species, how can these divergent results be reconciled?

### The role of the corticospinal tract

It must have felt uncanny to those early researchers to find that surface stimulation of the cortex produces discrete muscle responses, in a way so similar to what Galvani did with the frog’s leg. Indeed, Sherrington himself conveys the feeling clearly in the opening of his seminal lecture on the motor cortex (Sherrington, 1906, p.271), confessing “that although it is not surprising that such territorial subdivision of function should exist in the cerebral cortex, it is surprising that by our relatively imperfect artifices for stimulation we should be able to obtain clear evidence thereof.”

Of course, it did not go unnoticed that this fact might be due to the massive projection from cortex to the spinal cord, which had been fully traced by Ludwig Türck only twenty years before Fritsch and Hitzig’s experiment (Nathan and Smith, 1955). This corticospinal tract was found to originate in the anterior regions of the cerebral cortex and terminate directly in the lateral columns of the spinal cord after decussating (i.e. crossing over) at the level of the brainstem’s *medulla oblongata*. The existence of this corticospinal pathway presented compelling anatomical evidence of the means by which the motor cortex might be able to exert a direct influence on movement by electrical conduction of nerve impulses, but the role of this connection remained elusive.

### The effects of lesions in the corticospinal tract

In the wake of the Goltz-Ferrier debates, investigations of the role of the direct corticospinal descending pathway were conducted in multiple animal species. Sherrington himself started out his work by tracing spinal cord degeneration over large periods of time (up to 11 months) following cortical lesions in Goltz’s dogs (Langley and Sherrington, 1884; Sherrington, 1885). He confirmed that many of the properties of the corticospinal tract in the primate held for the dog, and furthermore became one of the first to observe the presence of a degenerated “re-crossed” pyramidal tract that travels down the cord ipsilateral to the side of the lesion (Sherrington, 1885). These fibers would later come to be called the ipsilateral, ventral corticospinal tract, and have since been found and described in most mammalian species as forming roughly 10% of the entire corticospinal projections (Kuypers, 1981; Brösamle and Schwab, 2000; Lacroix et al., 2004). However, he also had the chance during this time to observe first hand the negative effects of corticospinal degeneration following lesion, which had been previously reported by Goltz and others in a variety of non-primate specimens. In his own words:

> That the pyramidal tracts are in the dog requisite for volitional impulses to reach limbs and body seems negatived by the fact that the animal can run, leap, turn to either side, use neck and jaws, &c. with ease and success after nearly, if not wholly, complete degeneration of these tracts on both sides. Further, after complete degeneration of one pyramid, there is in the dog no obvious difference between the movements of the right and left sides. ((Sherrington, 1885, p.189))

Interestingly, he does note that “defect of motion is observable only as a clumsiness in execution of fine movements” (Sherrington, 1885). These observations once again stood out in stark contrast with lesion experiments reported by Ferrier in the monkey, where cauterization of specific motor cortical areas produced complete and persistent paralysis of the corresponding body parts (Ferrier and Yeo, 1884).

Years later, Sherrington would come back to the motor cortex with a new set of studies on stimulation and ablation of the precentral region (Grünbaum and Sherrington, 1903; Graham Brown and Sherrington, 1913; Leyton and Sherrington, 1917). In these studies together with Grünbaum, Sherrington targeted motor cortical lesions to the excitable area of the arm or the leg and tracked the recovery of the animals over time. Following the initial paresis and loss of muscle control they observed dramatic recovery of most skilled motor acts, such as peeling open a banana or climbing cages (Leyton and Sherrington, 1917). In order to test whether the recovery process was due to cortical reorganization, they systematically stimulated the areas adjacent to the lesion as well as the motor cortex of the opposite hemisphere, but failed to evoke movements in the affected limb (Leyton and Sherrington, 1917), as would be expected if commands were traveling down the corticospinal tract in spared regions. Furthermore, subsequent ablation of those areas failed to produce any new impairments in the recovered limb, leaving Sherrington and his colleagues at a loss to find the locus of recovery (Leyton and Sherrington, 1917).

Confused by these results, which they thought “caused concern to, students of cerebral physiology”, Glees and Cole introduced a set of more quantitative behavioural assays in the hope of tracking in detail the recovery of motor control (Glees and Cole, 1950; Cole, 1952). They studied the behaviour of monkeys solving various puzzle boxes following successive circumscribed lesions to the thumb, index and arm areas of the motor cortex. As Sherrington reported, there was a quick recovery after an initial period of paralysis and loss of motor control. However, even though the monkeys fully recovered their ability to skillfully open the puzzle box, some subtle movement deficits and paresis in the control of fine movements of the digits was reported to persist (Glees and Cole, 1950). When stimulating motor cortical areas surrounding the circumscribed lesions, they were able to evoke movements in the impacted digits and reinstate the paretic symptoms after further ablation (Glees and Cole, 1950). This suggested the hypothesis that surrounding areas of the motor cortex could undergo reorganization following the lesion. However, an important difference to emphasize between these experiments and those of Sherrington is the fact that only relatively circumscribed motor cortical regions were removed in each surgery, whereas in the original Sherrington study the entire elbow, wrist, index, thumb and remaining digit motor areas were excised at once (Leyton and Sherrington, 1917), most likely causing degeneration of the entire corticospinal pathway for the affected limb. The presence of an intact corticospinal tract, excitability of movements to low-current stimulation and transient paretic symptoms following ablation thus seem to go hand in hand.

In the hopes of clarifying the confusion of exactly which movements were controlled by cortex, other studies focused on lesions restricted to the corticospinal tract, using both unilateral and bilateral section at the level of the medullary pyramids (Tower, 1940; Lawrence and Kuypers, 1968a,b). The goal was to isolate the effects of all the individual descending pathways to the spinal cord and resolve once and for all the question of whether the corticospinal tract of the motor cortex was the source of all “voluntary” movements. Sarah Tower was the first to describe in detail the results of unilateral and bilateral pyramidotomy in primates, with and without lesion of the motor cortex (Tower, 1940). She summarized the condition as “hypotonic paresis”, characterized by a loss of skeletal muscle tone and depression of the vasomotor system, along with general weakening of the reflexes involving the affected limb segments. Although all discrete usage of the hand and digits was eliminated, she did emphasize the clear presence of voluntary movements in the various purposeful compensations produced by the animals to deal with the a[iction. Tower attributed these compensations to the preserved capacities of brainstem circuits.

A more definitive study to dissociate the effects of direct corticospinal and indirect brainstem descending pathways was conducted by Lawrence and Kuypers, and presented in their now classical publications (Lawrence and Kuypers, 1968a,b). Using the Klüver board, a task where monkeys have to pick morsels of food from differently sized round holes, they observed that while normal monkeys routinely pick up the food by pinching individual bits with their fingers, monkeys with bilateral corticospinal lesions were mostly unable to perform this precise pincer movement, and instead employed coarser compensatory clasping strategies to retrieve the food (Lawrence and Kuypers, 1968a). In addition, lesioned monkeys were consistently reported to be somewhat slower and less agile than normal animals. However, most of their overall movement repertoire was surprisingly preserved. Their final conclusions fit remarkably well with the initial observations of Sherrington in the dog, suggesting that the corticospinal pathways superimpose speed and agility on subcortical mechanisms, and provide the capacity for fractionation of movements such as independent finger movements (Lawrence and Kuypers, 1968a). These observations recapitulate the effects of motor cortical lesions reported by Sherrington, but remain at odds with the primary role assigned to motor cortex, and the direct corticospinal tract, with the control of all voluntary movements.

### There are anatomical differences in corticospinal projections between primates and other mammals

In primates, the conspicuous effects of motor cortical lesion can also be induced by sectioning the corticospinal tract, the direct monosynaptic projection that connects motor cortex, and other cortical regions, to the spinal cord (Tower, 1940; Lawrence and Kuypers, 1968a). In monkeys, and similarly in humans, this pathway has been found to directly terminate on spinal motor neurons responsible for the control of distal muscles (Leyton and Sherrington, 1917; Bernhard and Bohm, 1954) and is also thought to support the low-current movement responses evoked by electrical stimulation of the cortex, as evidenced by the increased difficulty in obtaining a stimulation response following section at the level of the medulla (Woolsey et al., 1972).

However, the corticospinal tract is by no means the only pathway from cortex to movement (Figure 1). Motor cortex targets many other brain regions that can themselves generate movement. In fact, this specialized connection from telencephalon to spinal cord appeared only recently in vertebrate evolution (ten Donkelaar, 2009), and was further elaborated to include a direct connection from cortex to motor neurons only in some primate species and other highly manipulative mammals such as raccoons (Heffner and Masterton, 1983). In all other mammals, including cats and rats, the termination pattern of the corticospinal tract largely avoids the motor neuron pools in ventral spinal cord and concentrates instead on intermediate zone interneurons and dorsal sensory neurons (Kuypers, 1981; Yang and Lemon, 2003). Why then is there such a large dependency on this tract for human motor control? One possibility is that the rubrospinal tract—a descending pathway originating in the brainstem and terminating in the intermediate zone—is degenerated in humans compared to other primates and mammals (Nathan and Smith, 1955, 1982), and is thought to play a role in compensating for the loss of the corticospinal tract in non-human species (Lawrence and Kuypers, 1968b; Zaaimi et al., 2012).

**Figure 1.**
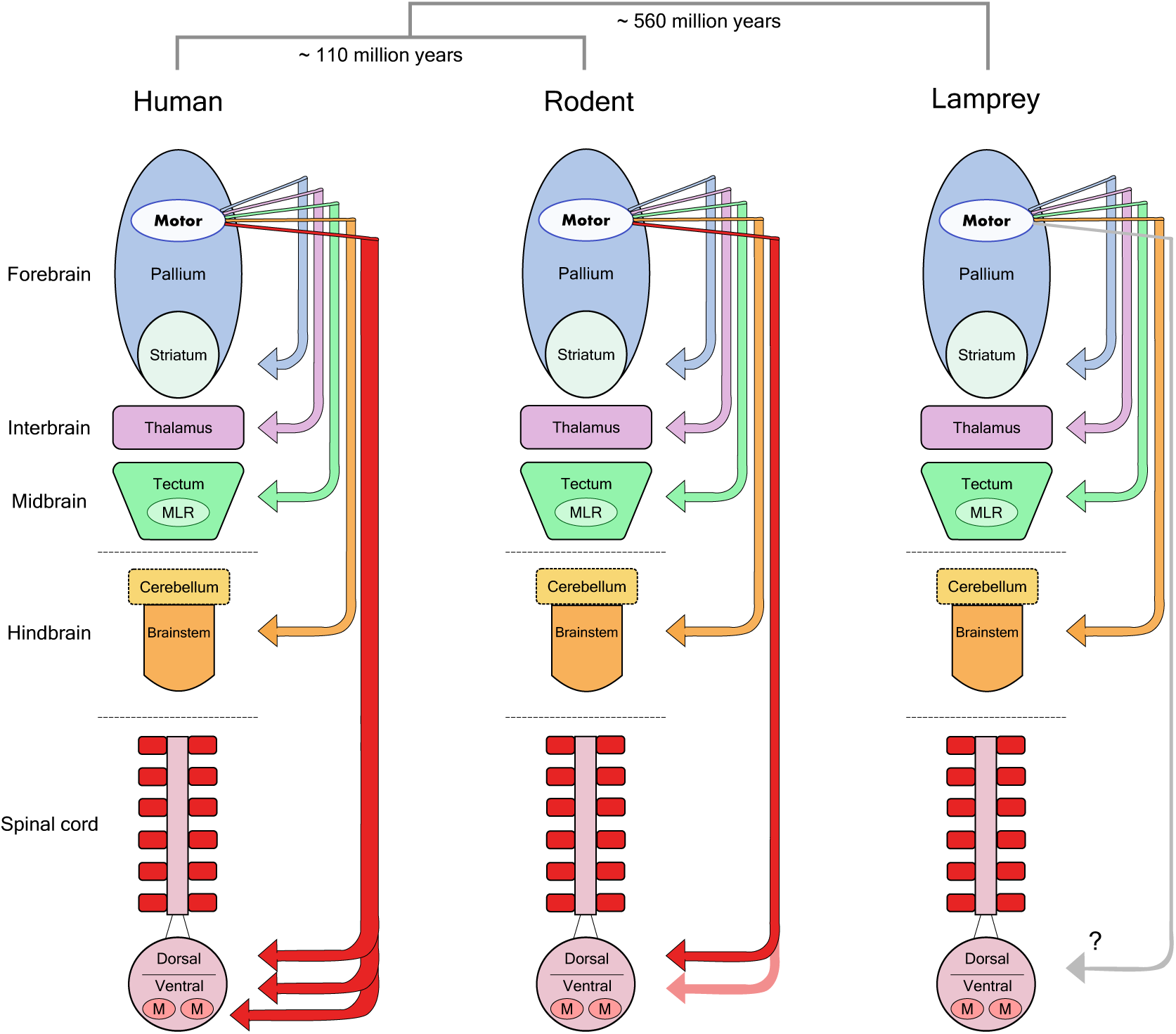
Forebrain motor control pathways across different vertebrate taxa. The molecular divergence times between human (primate), rodent and lamprey groups Kumar and Hedges (1998) are noted above a schematic view of the major divisions in the vertebrate brain. Arrows indicate the descending monosynaptic projections identified in each group from motor regions of the forebrain pallium to lower motor centres. Note the specialized monosynaptic projection directly targeting spinal motor neurons in human. MLR, Mesencephalic Locomotor Region; M, Motor Neurons.

It thus seems likely that most mammals rely on “indirect” pathways to convey cortical motor commands to muscles. These differences in anatomy might explain the lack of conspicuous, lasting movement deficits following motor cortical lesion in non-primates, but leaves behind a significant question: what is the motor cortex actually controlling in all these other mammals?

### What is the role of motor cortex in non-primate mammals?

In the rat, a large portion of cortex is considered “motor” based on anatomical (Donoghue and Wise, 1982), stimulation (Donoghue and Wise, 1982; Neafsey et al., 1986) and electrophysiological evidence (Hyland, 1998). However, the most consistently observed long-term motor control deficit following motor cortical lesion has been an impairment in supination of the wrist and individuation of digits during grasping, which in turn impairs reaching for food pellets through a narrow vertical slit (Whishaw et al., 1991; Alaverdashvili and Whishaw, 2008). Despite the fact that activity in rodent motor cortex has been correlated with movements in every part of the body (not just distal limbs) (Hill et al., 2011; Erlich et al., 2011), it would appear we are led to conclude that this large high-level motor structure, with dense efferent projections to motor areas in the spinal cord (Kuypers, 1981), basal ganglia (Turner and DeLong, 2000; Wu et al., 2009), thalamus (Lee et al., 2008), cerebellum (Baker et al., 2001) and brainstem (Jarratt and Hyland, 1999), as well as to most primary sensory areas (Petreanu et al., 2012; Schneider et al., 2014), evolved simply to facilitate more precise wrist rotations and grasping gestures. Maybe we are missing something. Might there be other problems in movement control that motor cortex is solving, but that we may be overlooking with our current assays?

### A role in modulating the movements generated by lower motor centres

The idea that the descending cortical pathways superimpose speed and precision on an existing baseline of behaviour has been suggested by lesion work in the primate (Lawrence and Kuypers, 1968b), but has been investigated much more thoroughly in the context of studies on the neural control of locomotion in cats. These studies have suggested that the corticospinal tract can play a role in the *adjustment* of ongoing movements, modulating the activity and sensory feedback in spinal circuits in order to adapt a lower movement controller to challenging conditions.

It has been known for more than a century that completely decerebrate cats are capable of sustaining the locomotor rhythms necessary for walking on a flat treadmill utilizing only spinal circuits (Graham Brown, 1911). In addition, there is a general capacity for spinal circuits to modulate network activity with incoming sensory input in order to coordinate and switch between different responses, even during specific phases of movement (Forssberg et al., 1975). Brainstem and mid-brain circuits are sufficient to initiate the activity of these spinal central pattern generators (Grillner and Shik, 1973), so what exactly is the contribution of motor cortex to the control of locomotion? Single-unit recordings of pyramidal tract neurons (PTNs) from cats walking on a treadmill have shown that a large proportion of these neurons are locked to the step cycle (Armstrong and Drew, 1984). However, we know from the decerebrate studies that this activity is not necessary for the basic locomotor pattern. What then is its role?

Lesions of the lateral descending pathways (containing corticospinal and rubrospinal projections) produce a long term impairment in the ability of cats to step over obstacles (Drew et al., 2002). Recordings of PTN neurons during locomotion show increased activity during these visually guided modifications to the basic step cycle (Drew et al., 1996). These observations suggest that motor cortex neurons are necessary for precise stepping and adjustment of ongoing locomotion to changing conditions. However, long-term effects seem to require complete lesion of *both* the corticospinal and rubrospinal tracts (Drew et al., 2002). Even in these animals, the voluntary act of stepping over an obstacle does not disappear entirely, and moreover, they can adapt to changes in the height of the obstacles (Drew et al., 2002). Although they never regain the ability to gracefully clear an obstacle, these animals still adjust their stepping height when faced with a higher obstacle in such a way that would have allowed them to comfortably clear the lower obstacle (Drew et al., 2002). Furthermore, deficits caused by lesions restricted to the pyramidal tract seem to disappear over time (Liddell and Phillips, 1944), and are most clearly visible only the first time an animal encounters a new obstacle (Liddell and Phillips, 1944).

The view that motor cortex in non-primate mammals is principally responsible for adjusting on-going movement patterns generated by lower brain structures is appealing. What is this modulation good for? What does it allow an animal to achieve? How can we assay its necessity?

### Towards a new teleology; new experiments required

It should now be clear that the involvement of motor cortex in the direct control of all “voluntary movement” is human-specific. There is a role for motor cortex across mammals in the control of precise movements of the extremities, especially those requiring individual movements of the fingers, but these effects are subtle in non-primate mammals. Furthermore, what would be a devastating impairment for humans may not be so severe for mammals that do not depend on precision finger movements for survival. Therefore, generalizing this specific role of motor cortex from humans to all other mammals would be misleading. We could be missing another, more primordial role for this structure that predominates in other mammals, and by doing so, we may also be missing an important role in humans.

The proposal that motor cortex induces modifications of ongoing movement synergies, prompted by the electrophysiological studies of cat locomotion, definitely points to a role consistent with the results of various lesion studies. However, in assays used thus far, the ability to modify ongoing movement generally recovers after a motor cortical lesion. What are the environmental situations in which motor cortical modulation is most useful?

Cortex has long been proposed to be the structure responsible for integrating a representation of the world and improving the predictive power of this representation with experience (Barlow, 1985; Doya, 1999). If motor cortex is the means by which these representations can gain influence over the body, however subtle and “modulatory”, can we find situations (i.e. tasks) in which this cortical control is required?

The necessity of cortex for various behavioural tasks has been actively investigated in experimental psychology for over a century, including the foundational work of Karl Lashley and his students (Lashley, 1921, 1950). In the rat, large cortical lesions were found to produce little to no impairment in movement control, and even deficits in learning and decision making abilities were difficult to demonstrate consistently over repeated trials. However, Lashley did notice some evidence that cortical control may be involved in postural adaptations to unexpected perturbations (Lashley, 1921). These studies once again seem to recapitulate the two most consistent observations found across the entire motor cortical lesion literature in non-primate mammals since Hitzig (Fritsch and Hitzig, 1870), Goltz (Goltz, 1888), Sherrington (Sherrington, 1885) and others (Oakley, 1979; Terry et al., 1989). One, direct voluntary control over movement is most definitely not abolished through lesion; and two, certain aspects of some movements are definitely impaired, but only under certain challenging situations. The latter are often reported only anecdotally. It was this collection of intriguing observations in animals with motor cortical lesions that prompted us to expand the scope of standard laboratory tasks to include a broader range of motor control challenges that brains encounter in their natural environments.

## Experiment Introduction

In the natural world, an animal must be able to adapt locomotion to any surface, not only in anticipation of upcoming terrain, but also in response to the unexpected perturbations that often occur during movement. This allows animals to move robustly through the world, even when navigating a changing environment. Testing the ability of the motor system to generate a robust response to an unexpected change can be difficult as it requires introducing a perturbation without cueing the animal about the altered state of the world. Marple-Horvat and colleagues built a circular ladder assay for cats that was specifically designed to record from motor cortex during such conditions (Marple-Horvat et al., 1993). One of the modifications they introduced was to make one of the rungs of the ladder fall unexpectedly under the weight of the animal. When they recorded from motor cortical neurons during the rung drop, they noticed a marked increase in activity, well above the recorded baseline from normal stepping, as the animal recovered from the fall and resumed walking. However, whether this increased activity of motor cortex was necessary for the recovery response has never been assayed.

#### Box 1. Some cautionary remarks on lesion techniques

The original methods used to induce a permanent lesion to the motor cortex were very crude, often involving gross mechanical aggression to the neural tissue by using surgical knife cuts or ablation by water-jet, aspiration, and thermo- or electrocoagulation. These methods are still widely used in lesion studies for their simplicity and bluntness, but have the disadvantage of making it hard to limit the lesion to a single area because of possible damage to subcortical areas or the destruction of fibers of passage. These limitations made it more difficult to interpret the effects of cortical lesions, and eventually led to the development of new techniques designed to work around such problems. Chemical injections of neurotoxic compounds such as ibotenic acid or kainic acid aim to increase selectivity of the lesion by limiting damage to neural cell bodies in the target area while leaving the fibers of passage intact (Schwarcz et al., 1979). Photothrombosis (Watson et al., 1985) or devascularization by pial stripping (Meyer and Meyer, 1971) aim to reproduce the effects of clinical stroke while avoiding extension of the lesion to subcortical areas as much as possible.

The early studies of Broca localizing the function of articulate language to a specific region in the cerebral hemispheres (Broca, 1861) established a long tradition of correlating the location of surgical brain injury with detailed analysis of any subsequent behavioural deficits. This method is not without its difficulties. The problems of plasticity and diaschisis will forever complicate conclusions based on injury and manipulation of nervous tissue (Lashley, 1933). Many recent methods for reversible chemical or optogenetic inactivation of the cortex have been proposed to improve statistical power of behavioural assessments (DeFeudis, 1980; Dong et al., 2010; Guo et al., 2015). Unfortunately, given that the cortex maintains a tight balance of excitation and inhibition during normal functioning and is also densely interconnected with the rest of the brain, the effects of such transient manipulations are prone to cause multiple downstream effects that can confound inferences about behavioural relevance (Otchy et al., 2015). In this respect, they are similar to stimulation experiments in that they are very useful in determining that two areas are connected in a circuit, but not necessarily what the connection means. Of course, permanent lesions themselves can induce plasticity changes in the function of downstream and upstream circuits. The expectation, however, is that such changes represent a homeostatically stable state of the system, allowing simultaneous investigation of the limits of recovery, as well as the kinds of problems for which a fully intact structure is definitely required.

## Results

To investigate whether the intact motor cortex is required for the robust control of movement in response to unexpected perturbations, we designed a reconfigurable dynamic obstacle course where individual steps can be made stable or unstable on a trial-by-trial basis (Figure 2, also see Methods). In this assay, rats shuttle back and forth across the obstacles, in the dark, in order to collect water rewards. We specifically designed the assay such that modifications to the physics of the obstacles could be made covertly. In this way, the animal has no explicit information about the state of the steps until it actually contacts them. Water deprived animals were trained daily for 4 weeks, throughout which they encountered increasingly challenging states of the obstacle course. Our goal was to characterize precisely the conditions under which motor cortex becomes necessary for the control of movement, and this motivated us to introduce an environment with graded levels of uncertainty.

**Figure 2.**
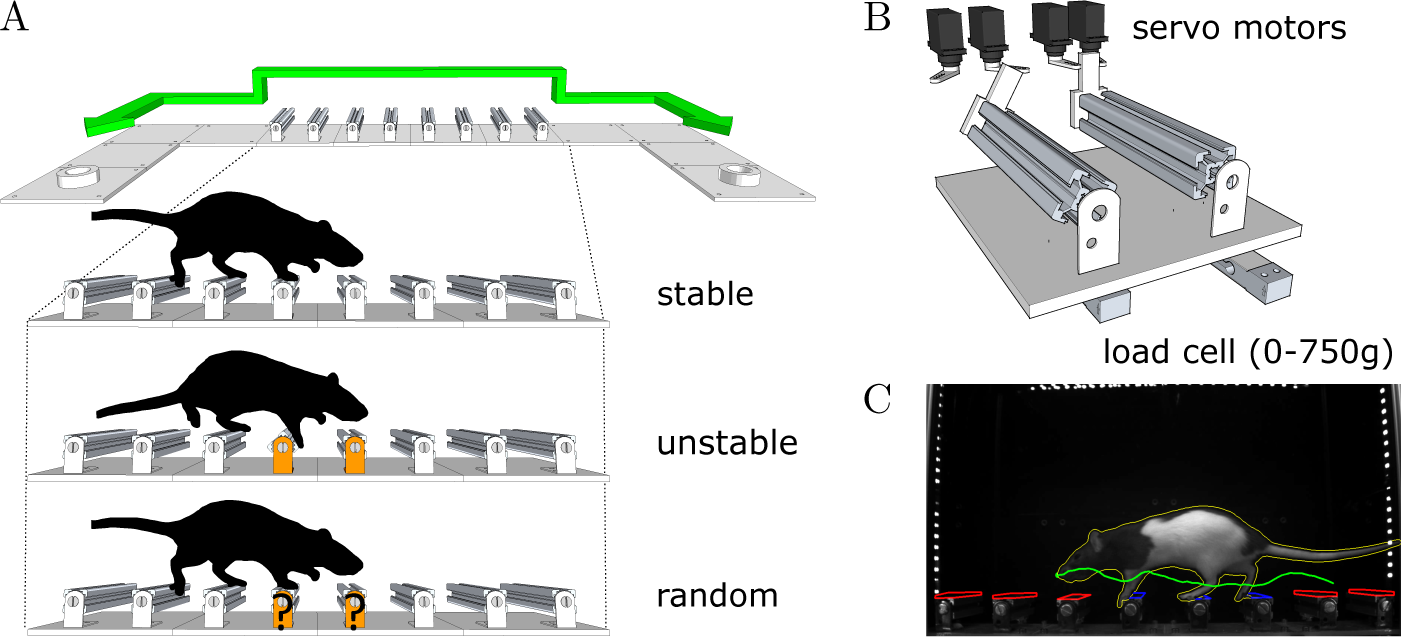
An obstacle course for rodents. (**A**) Schematic of the apparatus and summary of the different conditions in the behaviour protocol. Animals shuttle back and forth between two reward ports at either end of the enclosure. (**B**) Schematic of the locking mechanism that allows each individual step to be made stable or unstable on a trial-by-trial basis. (**C**) Example video frame from the behaviour tracking system. Coloured overlays represent regions of interest and feature traces extracted automatically from the video.

We compared the performance of 22 animals: 11 with bilateral ibotenic acid lesions to the primary and secondary forelimb motor cortex, and 11 age and gender matched controls (5 sham surgery, 6 wild-types). Animals were given ample time to recover, 4 weeks post-surgery, in order to specifically isolate behaviours that are chronically impaired in animals lacking the functions enabled by motor cortical structures. Histological examination of serial coronal sections revealed significant variability in the extent of damaged areas (Figure 3), which was likely caused by mechanical blockage of the injection pipette during lesion induction at some sites. Nevertheless, volume reconstruction of the serial sections allowed us to accurately quantify the size of each lesion, identify each animal (from Lesion A to Lesion K; largest to smallest), and use these values to compare observed behavioural effects as a function of lesion size.

**Figure 3.**
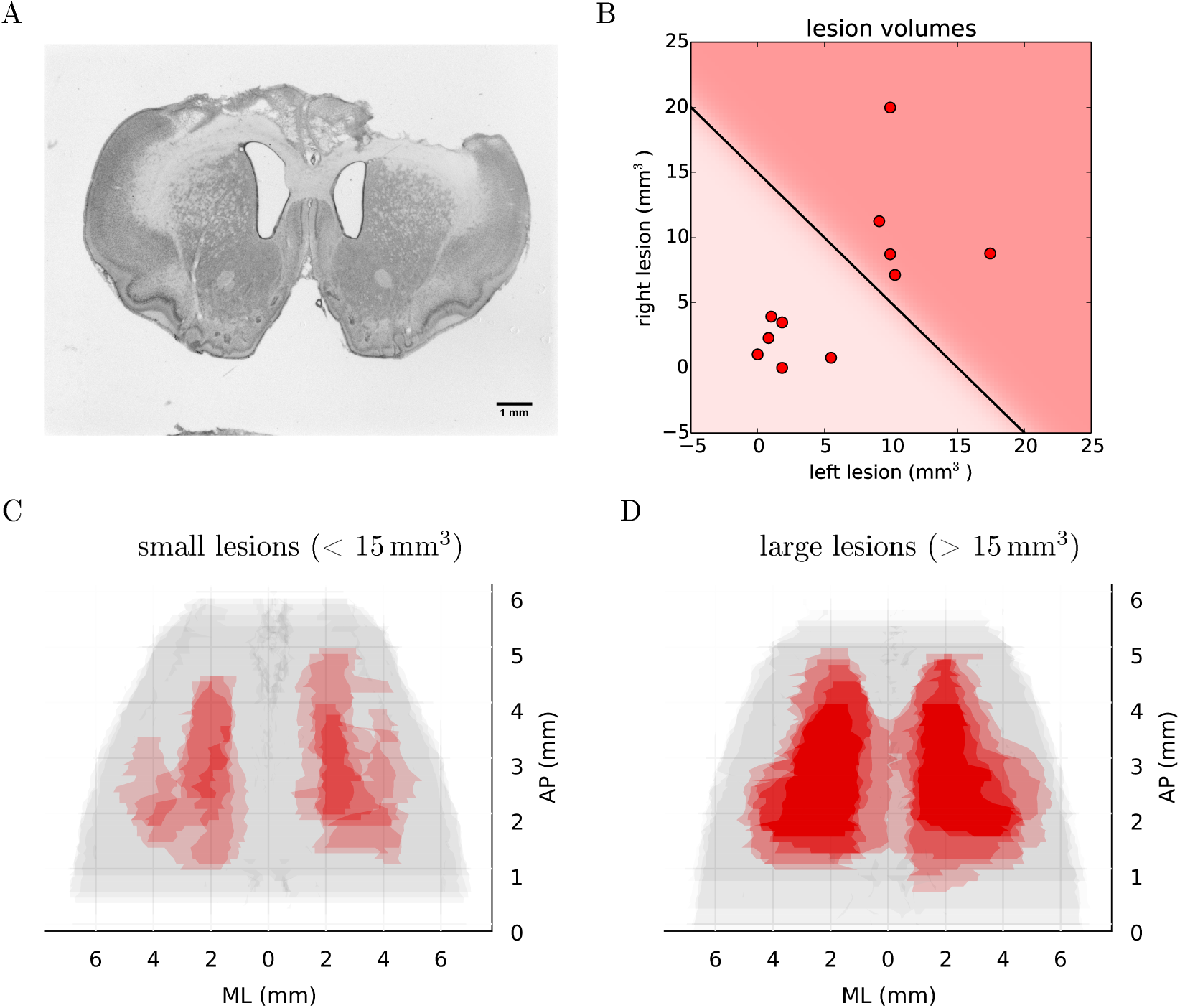
Histological analysis of lesion size. (**A**) Representative example of Nissl-stained coronal section showing bilateral ibotenic acid lesion of primary and secondary forelimb motor cortex. (**B**) Distribution of lesion volumes in the left and right hemispheres for individual animals. A lesion was considered “large” if the total lesion volume was above 15 mm^3^. (**C**) Super-imposed reconstruction stacks for all the small lesions (*n* = 6). (**D**) Super-imposed reconstruction stacks for all the large lesions (*n* = 5).

During the first sessions in the “stable” environment, all animals, both lesions and controls, quickly learned to shuttle across the obstacles, achieving stable, skilled performance after a few days of training (Figure 4). Even though the distance between steps was fixed for all animals, the time taken to adapt the crossing strategy was similar irrespective of body size. When first encountering the obstacles, animals adopted a cautious gait, investigating the location of the subsequent obstacle with their whiskers, stepping with the leading forepaw followed by a step to the same position with the trailing paw (Video 1: “First Leftwards Crossing”). However, over the course of only a few trials, all animals exhibited a new strategy of “stepping over” the planted forepaw to the next obstacle, suggesting an increased confidence in their movement strategy in this novel environment (Video 1: “Second Leftwards Crossing”). This more confident gait developed into a coordinated locomotion sequence after a few additional training sessions (Video 1: “Later Crossing”). The development of the ability to move confidently and quickly over the obstacle course was observed in both lesion and control animals (Video 2).

**Figure 4.**
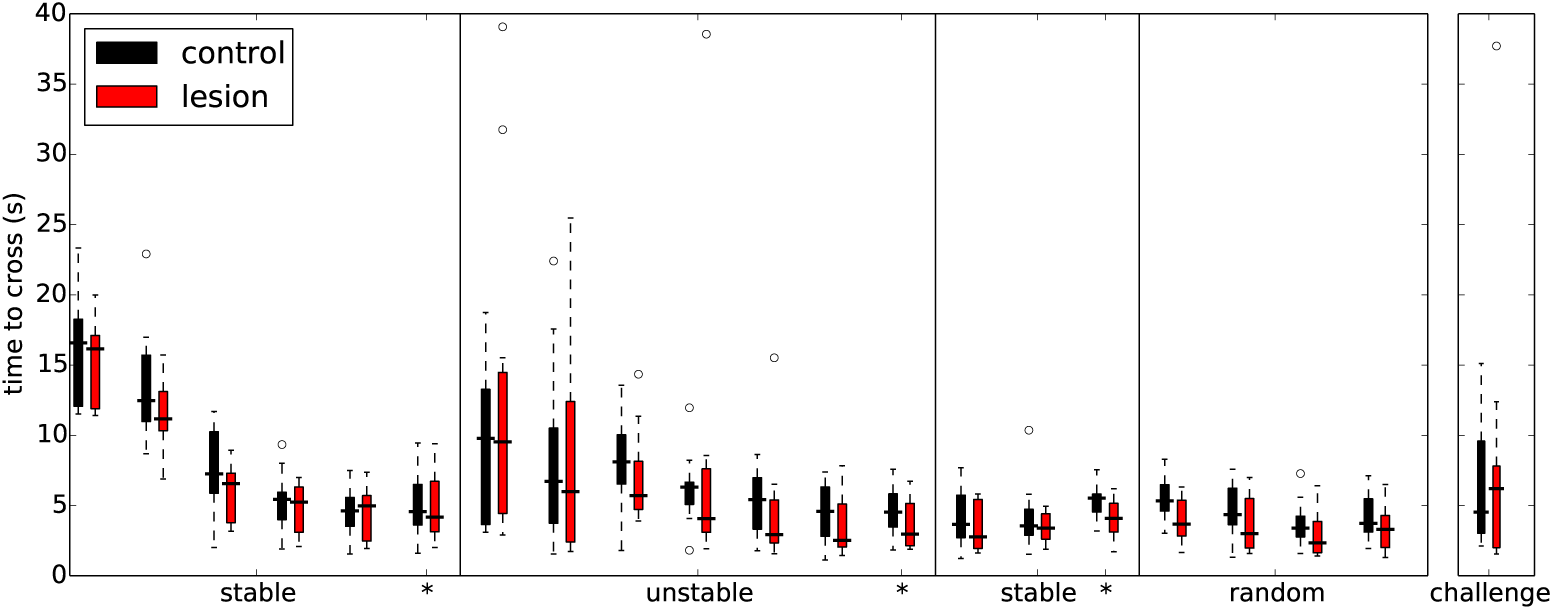
Overall performance on the obstacle course is similar for both lesion (*n* = 11) and control animals (*n* = 11) across the different protocol stages. Each set of coloured bars represents the distribution of average time to cross the obstacles on a single session. Asterisks indicate sessions where there was a change in assay conditions during the session (see text). In these transition sessions, the average performance on the 20 trials immediately preceding the change is shown to the left of the solid vertical line whereas the performance on the remainder of that session (after the change) is shown to the right.

In addition to the excitotoxic lesions, in three animals we performed larger frontal cortex aspiration lesions in order to determine whether the remaining trunk and hindlimb representations were necessary to navigate the elevated obstacle course. Also, in order to exclude the involvement of other corticospinal projecting regions in the parietal and rostral visual areas (Miller, 1987), we included three additional animals which underwent even more extensive cortical lesion procedures (Figure 5A,B, see Methods). These *extended* lesion animals were identified following chronological order (from Extended Lesion A to Extended Lesion F; where the first three animals correspond to frontal cortex aspiration lesions and the remaining animals to the more extensive frontoparietal lesions). In these extended cortical lesions, recovery was found to be overall slower than in lesions limited to the motor cortex, and animals required isolation and more extensive care during the recovery period.

Nevertheless, when tested in the shuttling assay, the basic performance of these extended lesion animals was similar to that of controls and animals with excitotoxic motor cortical lesions (Figure 5C). Animals with large frontoparietal lesions did exhibit a very noticeable deficit in paw placement throughout the early sessions (Figure 5D). Interestingly, detailed analysis of paw placement behaviour revealed that this deficit was almost entirely explained by impaired control of the hindlimbs. Paw slips were much more frequent when stepping with a hindlimb than with a forelimb (Figure 5E,F). In addition, when a slip did occur, these animals failed to adjust the affected paw to compensate for the fall (e.g. keeping their digits closed), which significantly impacted their overall posture recovery. These deficits in paw placement are consistent with results from sectioning the entire pyramidal tract in cats (Liddell and Phillips, 1944), and reports in ladder walking following motor cortical lesion in rodents (Metz and Whishaw, 2002), but surprisingly we did not observe deficits in paw placement in animals with ibotenic acid lesions limited to forelimb motor cortex (Figure 5D). Furthermore, despite this initial impairment, animals with extended lesions were still able to improve their motor control strategy up to the point where they were moving across the obstacles as efficiently as controls and other lesioned animals (Figure 5C, Video 2). Indeed, in the largest frontoparietal lesion, which extended all the way to rostral visual cortex, recovery of a stable locomotion pattern was evident over the course of just ten repeated trials (Video 3). The ability of this animal to improve its motor control strategy in such a short period of time seems to indicate the presence of motor learning, not simply an increase in confidence with the new environment.

**Figure 5.**
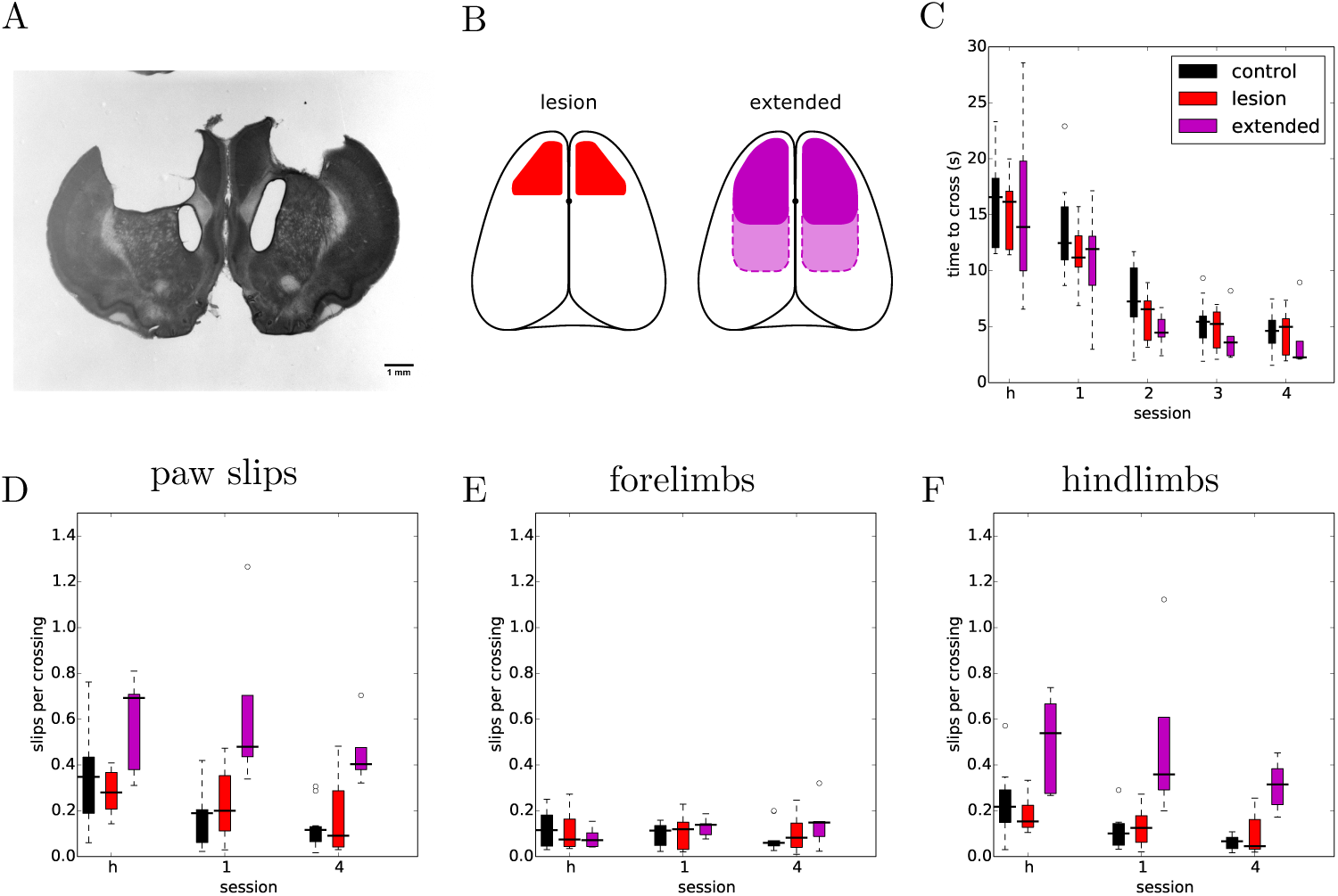
Extended frontoparietal cortex lesions perform as well as control animals despite impaired hindlimb control. (**A**) Representative example of Nissl-stained coronal section showing bilateral aspiration lesion of forelimb sensorimotor cortex. (**B**) Schematic depicting targeted lesion areas in the different animal groups. Left: outline of bilateral ibotenic acid lesions to the motor cortex. Right: outline of extended bilateral frontoparietal cortex lesions. Solid outline represents frontal cortex targeted lesions and dotted outline the more extensive frontoparietal lesions. (**C**) Average time required to cross the obstacles in the stable condition for extended lesions (*n* = 5). Performance of the other groups is shown for comparison. (**D**) Average number of slips per crossing in early versus late sessions of the stable condition. (**E**) Same data showing only forelimb slips. (**F**) Same data showing only hindlimb slips.

In subsequent training sessions we progressively increased the difficulty of the obstacle course, by making more steps unstable. The goal was to compare the performance of the two groups as a function of difficulty. Surprisingly, both lesion and control animals were able to improve their performance by the end of each training stage even for the most extreme condition where all steps were unstable (Figure 4, Video 4). This seems to indicate that the ability of these animals to fine-tune their motor performance in a challenging environment remained intact.

One noticeable exception was the animal with the largest ibotenic acid lesion. This animal, following exposure to the first unstable protocol, was unable to bring itself to cross the obstacle course (Video 5). Some other control and lesioned animals also experienced a similar form of distress following exposure to the unstable obstacles, but eventually all these animals managed to start crossing over the course of a single session. In order to test whether this was due to some kind of motor disability, we lowered the difficulty of the protocol for this one animal until it was able to cross again. Following a random permutation protocol, where any two single steps were released randomly, this animal was then able to cross a single released obstacle placed in any location of the assay. After this success, it eventually learned to cross the highest difficulty level in the assay in about the same time as all the other animals, suggesting that there was indeed no lasting motor execution or learning deficit, and that the disability must have been due to some other unknown, yet intriguing, (cognitive) factor.

Having established that the overall motor performance of these animals was similar across all conditions, we next asked whether there was any difference in the strategy used by the two groups of animals to cross the unstable obstacles. We noticed that during the first week of training, the posture of the animals when stepping on the obstacles changed significantly over time (Figure 6B,C). specifically, the centre of gravity of the body was shifted further forward and higher during later sessions, in a manner proportional to performance. However, after the obstacles changed to the unstable state, we observed an immediate and persistent adjustment of this crossing posture, with animals assuming a lower centre of gravity and reducing their speed as they approached the unstable obstacles (Figure 6C,D). Interestingly, we also noticed that a group of animals adopted a different strategy. Instead of lowering their centre of gravity, they either kept it unchanged or shifted it even more forward and performed a jump over the unstable obstacles (Figure 7A,B). These two strategies were remarkably consistent across the two groups, but there was no correlation between the strategy used and the degree of motor cortical lesion (Figure 6E,F, 7C). In fact, we found that the use of a jumping strategy was best predicted by the body weight of the animal (Figure 7C).

**Figure 6.**
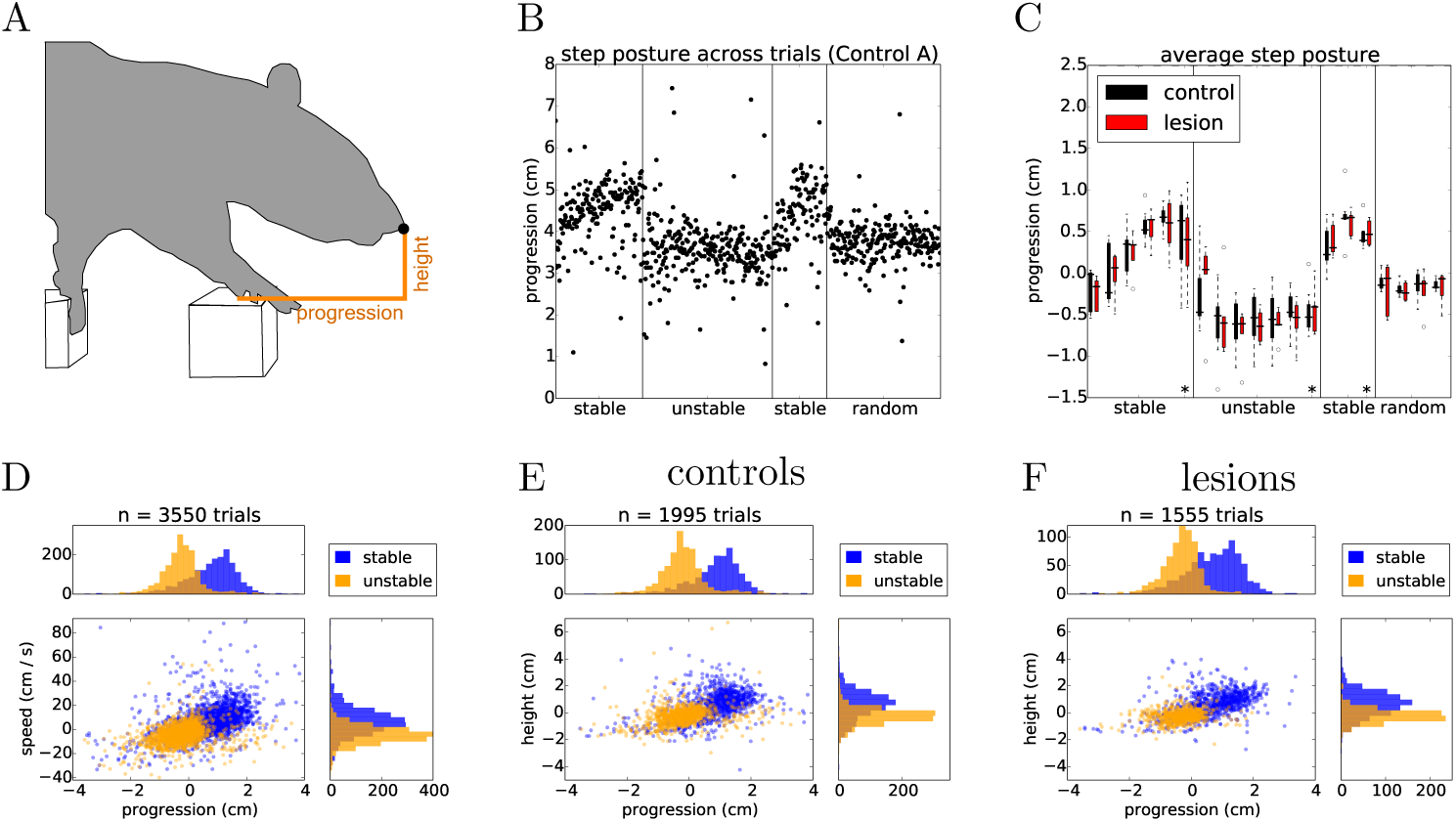
Rats adapt their postural approach to the obstacles after a change in physics. (**A**) Schematic of postural analysis image processing. The position of the animal’s nose is extracted whenever the paw activates the ROI of the first manipulated step (see methods). (**B**) The horizontal position, i.e. progression, of the nose in single trials for one of the control animals stepping across the different conditions of the shuttling protocol. (**C**) Average horizontal position of the nose across the different protocol stages for both lesion and control animals. Asterisks indicate the average nose position on the 20 trials immediately preceding a change in protocol conditions (see text). (**D**) Distribution of horizontal position against speed for the last two days of the stable (blue) and unstable (orange) protocol stages. (**E-F**) Distribution of nose positions for control and lesion animals over the same sessions.

**Figure 7.**
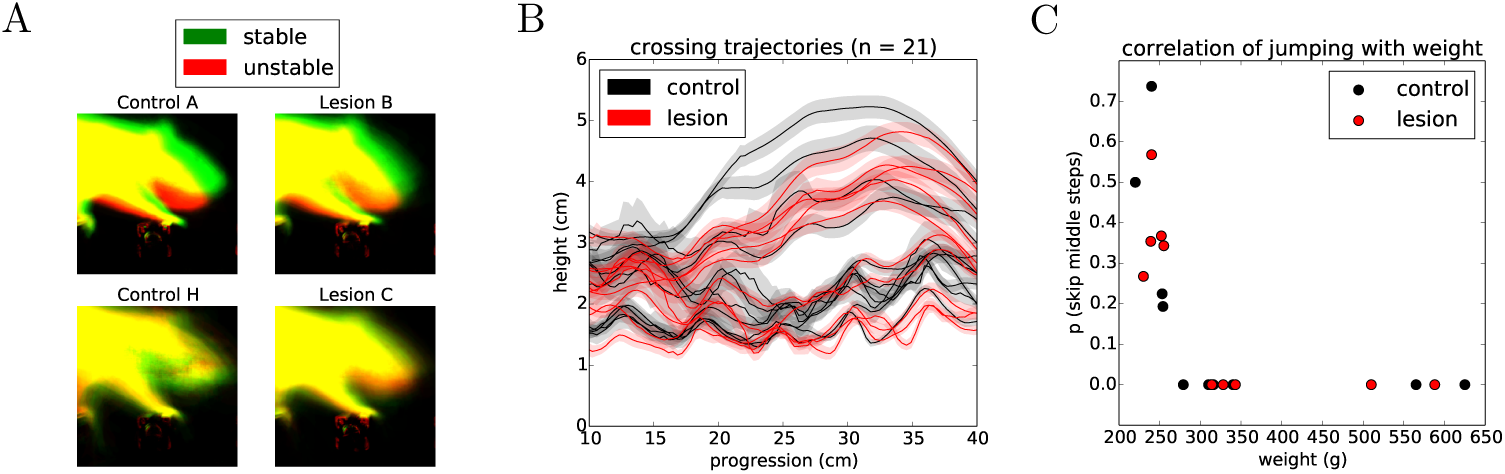
Animals use different strategies for dealing with the unstable obstacles. (**A**) Example average projection of all posture images for stable (green) and unstable (red) sessions for two non-jumper (top) and two jumper (bottom) animals. (**B**) Average nose trajectories for individual animals crossing the unstable condition. The shaded area around each line represents the 95% confidence interval. (**C**) Correlation of the probability of skipping the center two steps with the weight of the animal.

During the two days where the stable state of the environment was reinstated, the posture of the animals was gradually restored to pre-manipulation levels (Figure 6B,C), although in many cases this adjustment happened at a slower rate than the transition from stable to unstable. Again, this postural adaptation was independent of the presence or absence of forepaw motor cortex.

We next looked in detail at the days where the state of the obstacle course was randomized on a trial-by-trial basis. This stage of the protocol is particularly interesting as it reflects a situation where the environment has a persistent degree of uncertainty. For this analysis, we were forced to exclude the animals that employed a jumping strategy, as their experience with the manipulated obstacles was the same irrespective of the state of the world. First, we repeated the same posture analysis comparing all the stable and unstable trials in the random protocol in order to control for whether there was any subtle cue in our motorized setup that the animals might be using to gain information about the current state of the world. There was no significant difference between randomly presented stable and unstable trials on the approach posture of the animal (Figure 8A). However, classifying the trials on the basis of past trial history revealed a significant effect on posture (Figure 8B). This suggested that the animals were adjusting their body posture when stepping on the affected obstacles on the basis of their current expectation about the state of the world, which is updated by the previously experienced state. Surprisingly, this effect again did not depend on the presence or absence of frontal motor cortical structures (Figure 8C,D).

**Figure 8.**
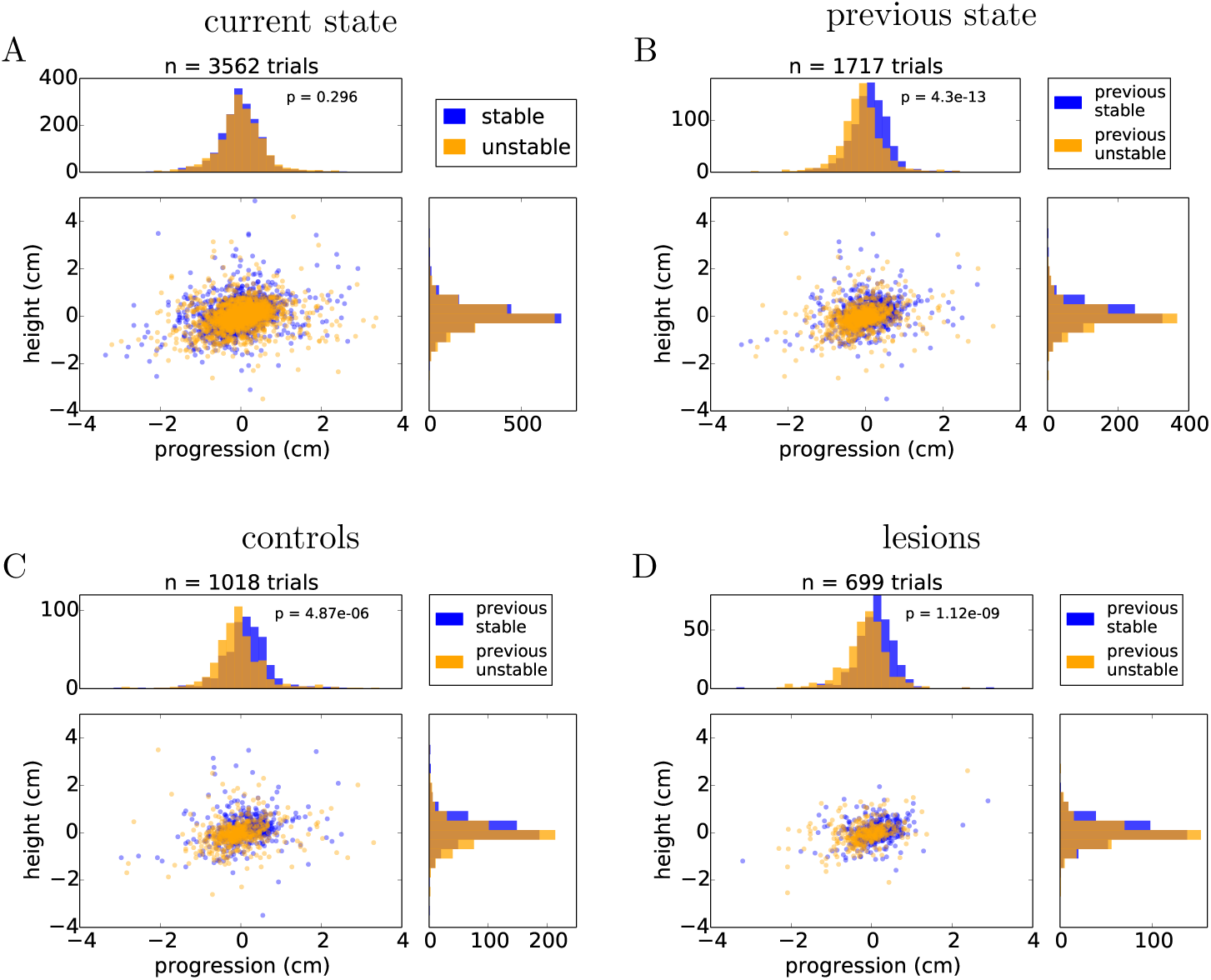
Animals adjust their posture on a trial-by-trial basis to the expected state of the world. (**A**) Distribution of nose positions on the randomized protocol when stepping on the first manipulated obstacle, for trials in which the current state was stable (blue) or unstable (orange). (**B**) Distribution of nose positions for trials in which the previous two trials were stable (blue) or unstable (orange). (**C-D**) Same data as in (**B**) split by the control and lesion groups. *p* values from Student’s unpaired t-test are indicated.

Finally, we decided to test whether general motor performance was affected by the randomized state of the obstacles. If the animals do not know what state the world will be in, then there will be an increased challenge to their stability when they cross over the unstable obstacles, possibly demanding a quick change in strategy when they learn whether the world is stable or unstable. In order to evaluate the dynamics of crossing, we compared the speed profile of each animal across these different conditions (Figure 9, see Methods). Interestingly, two of the animals with the largest lesions appeared to be significantly slowed down on unstable trials, while controls and the animals with the smallest lesions instead tended to accelerate after encountering an unstable obstacle. However, the overall effect for lesions versus controls was not statistically significant (Figure 9C).

**Figure 9.**
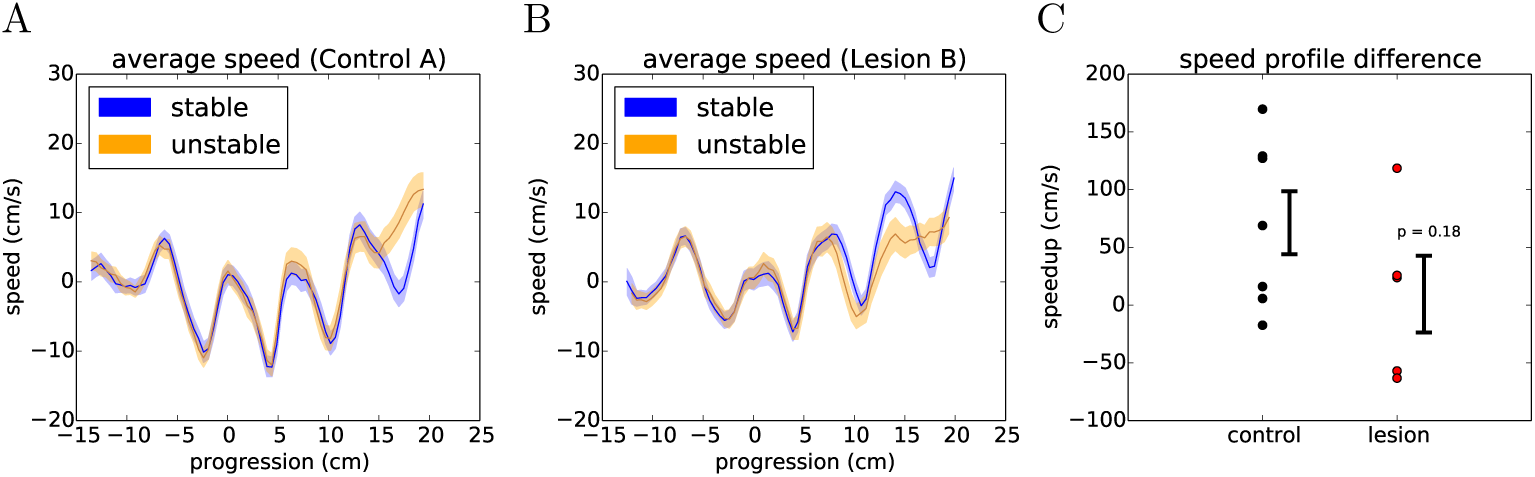
Encountering different states of the randomized obstacles causes the animals to quickly adjust their movement trajectory. (**A**) Example average speed profile across the obstacles for stable (blue) and unstable (orange) trials in the randomized sessions of a control animal (see text). The shaded area around each line represents the 95% confidence interval. (**B**) Respectively for one of the largest lesions. (**C**) Summary of the average difference between the speed profiles for stable and unstable trials across the two groups of animals. Error bars show standard error of the mean. *p* value from Student’s unpaired t-test is indicated.

Nevertheless, we were intrigued by this observation and decided to investigate, in detail, the first moment in the assay when a perturbation is encountered. In the random protocol, even though the state of the world is unpredictable, the animals know that the obstacles might become unstable. However, the very first time the environment becomes unstable, the collapse of the obstacles is completely unexpected and demands an entirely novel motor response.

A detailed analysis of the responses to the first collapse of the steps revealed a striking difference in the strategies deployed by the lesion and control animals. Upon the first encounter with the manipulated steps, we observed three types of behavioural responses from the animals (Video 6): investigation, in which the animals immediately stop their progression and orient towards, whisk, and physically manipulate the altered obstacle; compensation, in which the animals rapidly adjust their behaviour to negotiate the unexpected instability; and halting, in which the ongoing motor program ceases and the animals’ behaviour simply comes to a stop for several seconds. Remarkably, these responses depended on the presence or absence of motor cortex (Figure 10). Animals with the largest motor cortical lesions, upon their first encounter with the novel environmental obstacle, halted for several seconds, whereas animals with an intact motor cortex, and those with the smallest lesions, were able to rapidly react with either an investigatory or compensatory response (Video 7,8).

**Figure 10.**
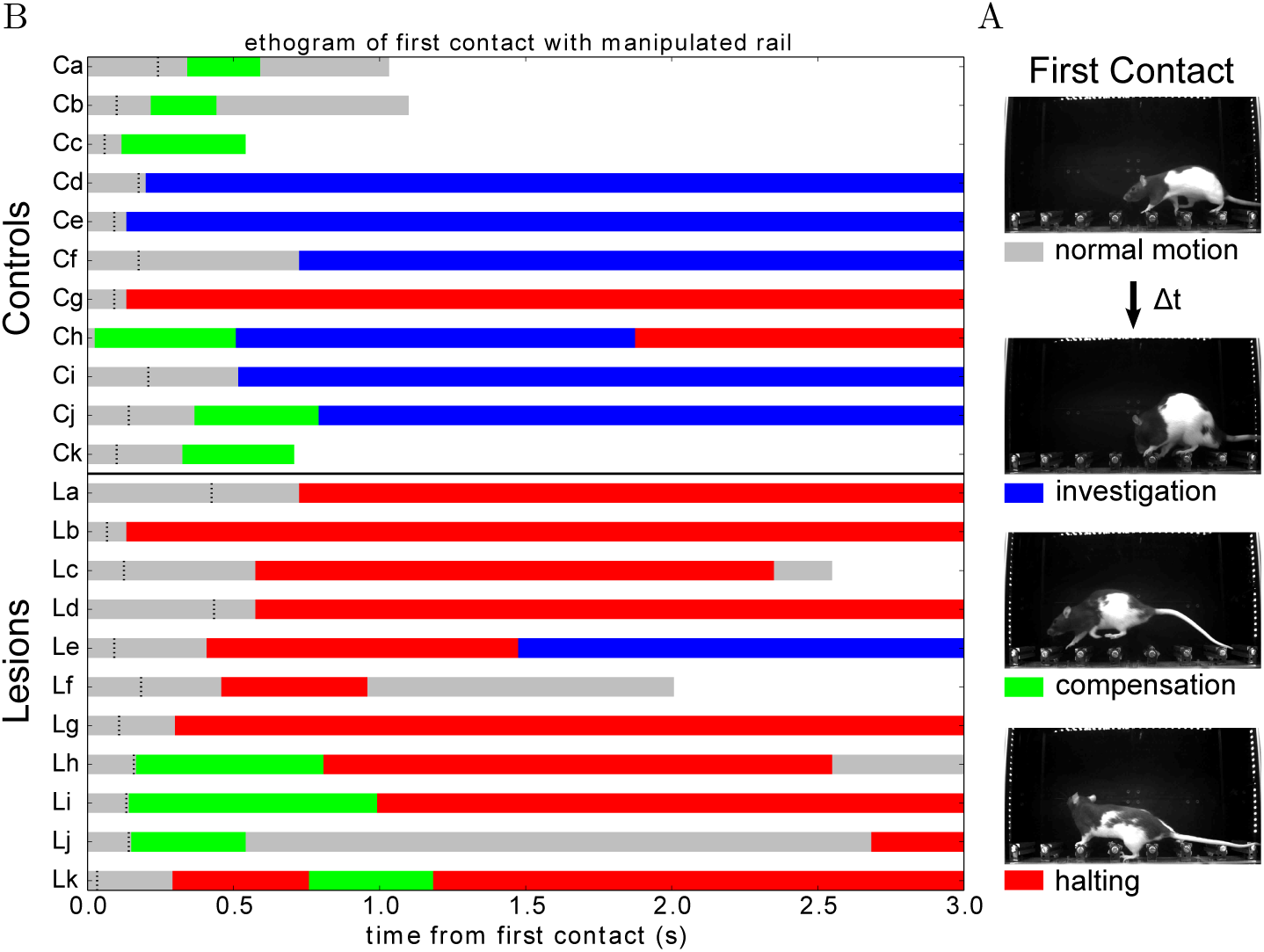
Responses to an unexpected change in the environment. (**A**) Response types observed across individuals upon first encountering an unpredicted instability in the state of the centre obstacles. (**B**) Ethogram of behavioural responses classified according to the three criteria described in (**A**) and aligned (0.0) on first contact with the newly manipulated obstacle. Black dashes indicate when the animal exhibits a pronounced ear flick. White indicates that the animal has crossed the obstacle course.

The response of animals with extended lesions was even more striking. In two of these animals, there was a failure to recognize that a change had occurred at all (Video 9). Instead, they kept walking across the now unstable steps for several trials, never stopping to assess the new situation. One of them gradually noticed the manipulation and stopped his progression, while the other one only fully realized the change after inadvertently hitting the steps with its snout (Video 9: Extended Lesion A). This was the first time we ever observed this behaviour, as all animals with or without cortical lesions always displayed a clear switch in behavioural state following the first encounter with the manipulation. In the remaining animals with extended lesions, two of them clearly halted their progression following the collapse of the obstacles, in a way similar to the large motor cortex ibotenic lesions (Video 10). The third animal (Extended Lesion B) actually collapsed upon contact with the manipulated step, falling over its paw and digits awkwardly and hitting the obstacles with its snout. Shortly after this there was a switch to an exploratory behaviour state, in a way similar to Extended Lesion A.

In order to investigate the neurophysiological correlates of these robust responses in the motor cortex, in three animals we implanted flexible surface electrode grids above the dura in one hemisphere of the intact brain (Figure 11A, also see Methods). Each step of the obstacle course was outfitted with a load cell sensor to measure the precise timing of contact and the amount of weight placed on each limb during locomotion. The entire electrocorticography (ECoG) system was synchronized on a frame-by-frame basis with the high-speed video acquisition so we could reconstruct the detailed behaviour of the animal at any point of the physiological trace as well as relate the continuous load profile on individual steps with different phases in the locomotion cycle.

**Figure 11.**
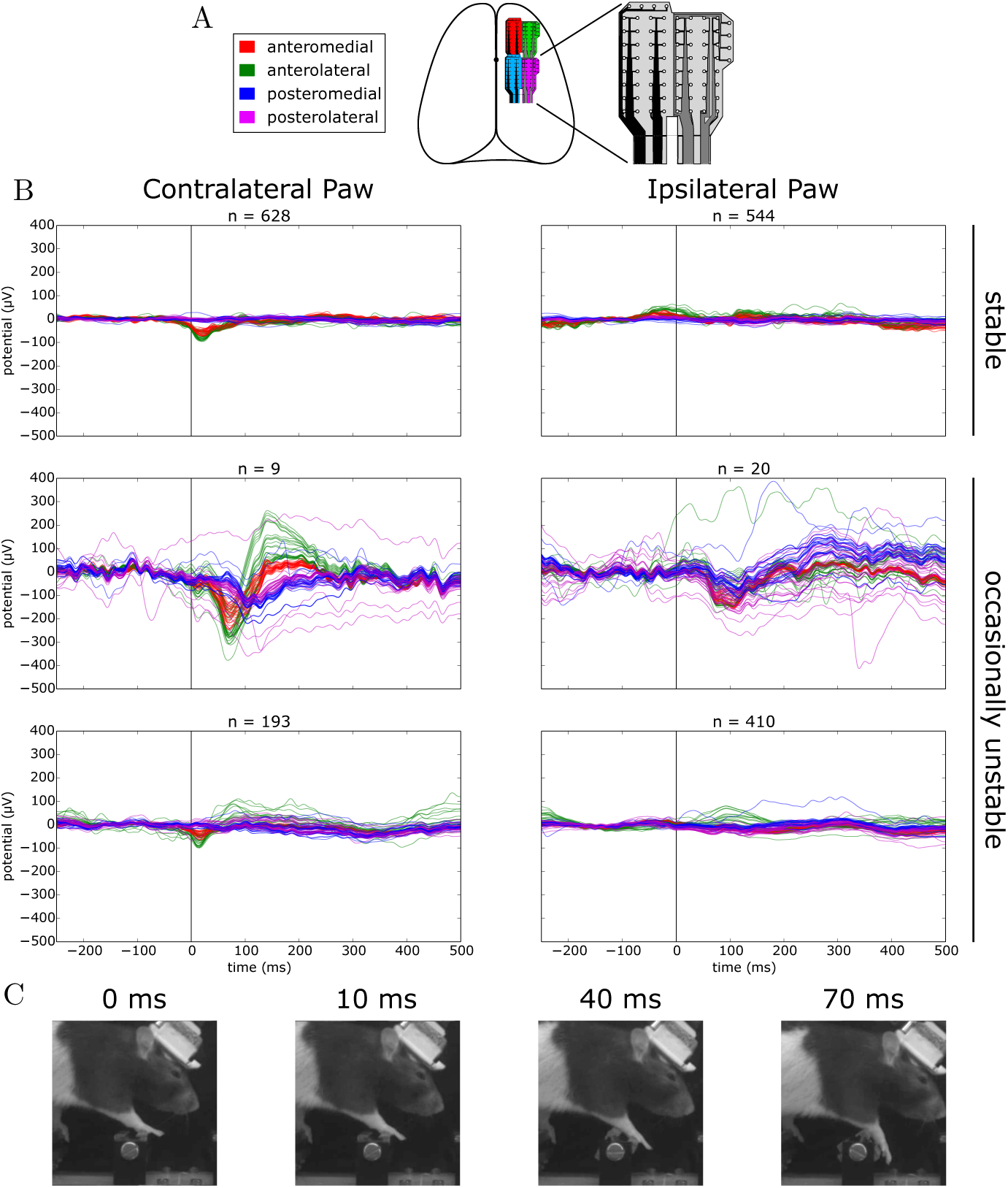
Evoked responses to stepping on stable versus occasionally unstable steps. (**A**) Schematic depicting the location of implanted ECoG grids. (**B**) Average voltage traces aligned on stepping with the contra- or ipsilateral paw on a manipulated step. Top: sessions where the step was permanently stable. Bottom: sessions where the step was occasionally made unstable. The middle row shows traces for unstable trials and the lower row the traces for the remaining stable trials. (**C**) Example frames of the behaviour of the animal at different time points of an unstable trial.

We first asked whether there were responses in the ECoG signal over forelimb motor cortex that were modulated by stepping behaviour. Aligning the ECoG traces to the event of stepping on a permanently stable step with the contralateral paw revealed the distinct presence of an evoked potential on the anterior grid channels that was absent when stepping with the ipsilateral paw (Figure 11B, top trace). On close inspection, it could be seen that the beginning of the negative deflection slightly precedes the time of contact with the step, suggesting a non-sensory contribution to the evoked response. Synaptic activity in the long and thick apical dendrites of pyramidal cells are thought to be one of the main contributors to cortically recorded extracellular field potentials (Buzsáki et al., 2012). In the cat, a sizeable proportion of pyramidal tract neurons in the motor cortex have been found to discharge rhythmically during unimpeded locomotion (Armstrong and Drew, 1984; Drew et al., 1996), a phenomenon that is very likely to be coupled with observable synaptic activity in the potential traces and could account for the step-aligned evoked responses that we observed during locomotion of rats in the stable obstacle course.

Next, we asked whether there was any modulation of the evoked response when navigating the unstable obstacle course. In order to try and maximize the number of trials in which the encounter with the unstable step is unexpected, we adjusted the behavioural protocol at the transition between the stable and unstable test periods. This time, instead of permanently switching the centre steps to the unstable configuration, we decided to immediately revert the steps back to the stable state after the first exposure to the instability. After 20 subsequent trials in the stable state, the steps were again made unstable, and this pattern was repeated for several days.

Surprisingly, when we aligned the ECoG traces to contralateral paw steps on the manipulated obstacle in unstable trials, we observed a second evoked negativity, delayed in time relative to the previously observed stable step evoked response, and with a much larger amplitude across the channels in the anterior grid (Figure 11B, middle left trace). Remarkably, even in the presence of such a small number of trials, the consistency of the response in every trial provided a sufficient signal-to-noise ratio for the average response to be clearly visible. Interestingly, this negativity was found to be rapidly followed by an equally large positive deflection in the potential which decayed to baseline with a much larger time constant, a response that was entirely absent from the evoked potential to stepping on a stable step. In contrast, the response to unstable steps with the ipsilateral paw did not reveal such large deflections from the baseline, although a consistent negativity could still be seen across the grid around the same time point (Figure 11B, middle right trace). The amplitude and timing of evoked responses when stepping with the contralateral paw on the same manipulated step in stable trials was largely identical to the condition where the step was permanently stable, and again was found to be absent when stepping with the ipsilateral paw (Figure 11B, bottom trace).

To investigate whether such a large evoked response correlated with an equally pronounced change in the overt behaviour of the animal, we extracted successive frames in the high-speed video corresponding to different time points of the trace (Figure 11C). Interestingly, there was no obvious motor response from the animal up to the point where the negativity peaks at around 70 ms. In fact, the affected paw was seen to mostly follow the inertia of the rotating step and no further motor response was observed before 100 ms, roughly consistent with the compensation reaction times observed in the responses to an unexpected collapse of the steps in control animals (Figure 10). The basic features of these evoked potential profiles were recapitulated across all the remaining animals (data not shown).

## Experiment Discussion

In this experiment, we assessed the role of motor cortical structures by making targeted lesions to areas implicated with forelimb control in the rat (Kawai et al., 2015; Otchy et al., 2015). Consistent with previous studies, we did not observe any conspicuous deficits in movement execution for rats with bilateral motor cortex lesions when negotiating a stable environment. Even when exposed to a sequence of unstable obstacles, animals were able to learn an efficient strategy for crossing these more challenging environments, with or without motor cortex. These movement strategies also include a preparatory component that might reflect the state of the world an animal expected to encounter. Surprisingly, these preparatory responses also did not require the presence of motor cortex.

It was only when the environment did not conform to expectation, and demanded a rapid adjustment, that a difference between the lesion and control groups was obvious. Animals with extensive damage to the motor cortex did not deploy a change in strategy. Rather, they halted their progression for several seconds, unable to robustly respond to the new motor challenge. In a natural setting, such hesitation could easily prove fatal. Control animals, on the other hand, were able to rapidly and flexibly reorganize their motor response to an entirely unexpected change in the environment.

Our preliminary investigations of the neurophysiological basis of these robust responses with ECoG have revealed the presence of large amplitude evoked potentials in the motor cortex arising specifically in response to an unexpected collapse of the steps during locomotion. Compared with evoked responses obtained from normal stepping under stable conditions (−100 µV peak at 10 ms), these potentials are both much larger (−300 µV) and delayed in time (peak at 70 ms). Still, they preceded any overt behaviour corrections from the animal following the perturbation, as observed in the high-speed video recordings. The onset of these evoked potentials is in the range of the long-latency stretch reflex, which has been suggested to involve a transcortical loop through the motor cortex (Phillips, 1969; Matthews et al., 1990; Capaday et al., 1991). However, the simultaneous complexity and rapidity of adaptive motor responses we observed in control animals is striking, as they appear to go beyond simple corrective responses to reach a predetermined goal and include a fast switch to entirely different investigatory or compensatory motor strategies adapted to the novel situation. What is the nature of these robust responses that animals without motor cortex seem unable to deploy? What do they allow an animal to achieve? Why are cortical structures necessary for their successful and rapid deployment?

## Extended Discussion

Is “robust control” a problem worthy of high level cortical input? Recovering from a perturbation, to maintain balance or minimize the impact of a fall, is a role normally assigned to our lower level postural control systems. The corrective responses embedded in our spinal cord (Sherrington, 1893, 1910), brainstem (Arshian et al., 2014) and midbrain (Grillner and Shik, 1973) are clearly important components of this stabilizing network, but are they sufficient to maintain robust movement in the dynamic environments that we encounter on a daily basis? Some insight into the requirements for a robust control system can be gained from engineering attempts to build robots that navigate in natural environments.

In the field of robotics, feats of precision and fine movement control (the most commonly prescribed role for motor cortex), are not a major source of difficulty. Industrial robots have long since exceeded human performance in both accuracy and execution speed (Senoo et al., 2009). More recently, using reinforcement learning methods, they are now able to automatically learn efficient movement strategies, given a human-defined goal and many repeated trials for fine-tuning (Coates et al., 2008). What then are the hard problems in robotic motor control? Why are most robots still confined to factories, i.e. controlled, predictable environments? The reason is that as soon as a robot encounters natural terrain, a vast number of previously unknown situations arise. The resulting “perturbations” are dealt with poorly by the statistical machine learning models that are currently used to train robots in controlled settings.

Let’s consider a familiar example: You are up early on a Sunday morning and head outside to collect the newspaper. It is cold out, so you put on a robe and some slippers, open the front door, and descend the steps leading down to the street in front of your house. Unbeknownst to you, a thin layer of ice has formed overnight and your foot is now quickly sliding out from underneath you. You are about to fall. What do you do? Well, this depends. Is there a railing you can grab to catch yourself? Were you carrying a cup of coffee? Did you notice the frost on the lawn and step cautiously, anticipating a slippery surface? Avoiding a dangerous fall, or recovering gracefully, requires a rich knowledge of the world, knowledge that is not immediately available to spinal or even brainstem circuits. This rich context relevant for robust movement is readily available in cortex, and cortex alone.

Imagine now that you are tasked with building a robot to collect your morning newspaper. This robot, in order to avoid a catastrophic and costly failure, would need to have all of this contextual knowledge as well. It would need to know about the structure of the local environment (e.g. hand railings that can support its weight), hot liquids and their viscosities, and even the correlation of frozen dew with icy surfaces. To be a truly robust movement machine, a robot must *understand* the physical structure of the world.

Reaching to stop a fall while holding a cup of coffee is not exactly the kind of feat for which we praise our athletes and sports champions, and this might explain why the difficulty of such “feats of robustness” are often overlooked. However, it would not be the first time that we find ourselves humbled by the daunting complexity of a problem that we naively assumed was trivial. Vision, for example, has remained an impressively hard task for a machine to solve at human-level performance, yet it was originally proposed as an undergraduate summer project (Papert, 1966). Perhaps a similar misestimate has clouded our designation of the hard motor control problems worthy of cortical input.

Inspired by the challenges confronting roboticists, as well as our rodent behavioural results, we are now in a position to posit a new role for motor cortex.

### A primordial role for motor cortex

We are seeking a role for motor cortex in non-primate mammals, animals that do not require this structure for overt movement production. The struggles of roboticists highlight the difficulty of building movement systems that robustly adapt to unexpected perturbations, and the results we report in this study suggest that this is, indeed, the most conspicuous deficit for rats lacking motor cortex. So let us propose that, in rodents, motor cortex is primarily responsible for extending the robustness of the subcortical movement systems. It is not required for control in stable, predictable, non-perturbing environments, but instead specifically exerts its influence when unexpected challenges arise. This, we propose, was the original selective pressure for evolving a motor cortex, and thus, its primordial role. This role persists in all mammals, mediated via a modulation of the subcortical motor system (as is emphasized in studies of cat locomotion), and has evolved in primates to include direct control of the skeletal musculature. Our proposal of a “robust” teleology for motor cortex has a number of interesting implications.

### Implications for non-primate mammals

One of the most impressive traits of mammals is the vast range of environmental niches that they occupy. While most other animals adapt to change over evolutionary time scales, mammals excel in their flexibility, quickly evaluating and responding to unexpected situations, and taking risks even when faced with challenges that have never been previously encountered (Spinka et al., 2001). This success requires more than precision, it requires resourcefulness: the ability to quickly come up with a motor solution for any situation and under any condition (Bernstein, 1996). The Russian neurophysiologist Bernstein referred to this ability with an unconventional definition of “dexterity”, which he considered to be distinct from a simple harmony and precision of movements. In his words, dexterity is required only when there is “a conglomerate of unexpected, unique complications in the external situations, [such as] in a quick succession of motor tasks that are all unlike each other” (Bernstein, 1996).

If Bernstein’s “robust dexterity” is the primary role for motor cortex, then it becomes clear why the effects of lesions have thus far been so hard to characterize: assays of motor behaviour typically evaluate situations that are repeated over many trials in a stable environment. Such repeated tasks were useful, as they offer improved statistical power for quantification and comparison. However, we propose that these conditions specifically exclude the scenarios for which motor cortex originally evolved. It is not easy to repeatedly produce conditions that animals have not previously encountered, and the challenges in analysing these unique situations are considerable.

The assay reported here represents our first attempt at such an experiment, and it has already revealed that such conditions may indeed be necessary to isolate the role of motor cortex in rodents. We thus propose that neuroscience should pursue similar assays, emphasizing unexpected perturbations and novel challenges, and we have developed new hardware and software tools to make their design and implementation much easier (Lopes et al., 2015).

### Implications for primate studies

In contrast to other mammals, primates require motor cortex for the direct control of movement. However, do they also retain its role in generating robust responses? The general paresis, or even paralysis, that results from motor cortical lesions in these species obscures the involvement of cortex in directing rapid responses to perturbations. Yet there is evidence that a role in robust control is still present in primates, including humans. For example, stroke patients with partial lesions to the distributed motor cortical system will often recover the ability to move the affected musculature. However, even after recovering movement, stroke patients are still prone to severe impairments in robust control: unsupported falls are one of the leading causes of injury and death in patients surviving motor cortical stroke (Jacobs, 2014). We thus suggest that stroke therapy, currently focused on regaining direct movement control, should also consider strategies for improving robust responses.

Even if we acknowledge that a primordial role of motor cortex is still apparent in primate movement control, it remains to be explained why the motor cortex of these species acquired direct control of basic movements in the first place. This is an open question.

### Some speculation on the role of direct cortical control

What happens when cortex acquires direct control of movement? First, it must learn how to use this influence, bypassing or modifying lower movement controllers. While functional corticospinal tract connections may be established prenatally (Eyre et al., 2000), the refinement of corticospinal dependent movements, which must override the lower motor system, takes much longer and coincides with the lengthy maturation period of corticospinal termination patterns (Lawrence and Hopkins, 1976). Humans require years of practice to produce and refine basic locomotion and grasping (Thelen, 1985; von Hofsten, 1989), motor behaviours that are available to other mammals almost immediately after birth. This may be the cost of giving cortex direct control of movement—it takes more time to figure out how to move the body—but what is the benefit?

Giving motor cortex direct control over the detailed dynamics of movement might simply have extended the range and flexibility of robust responses. This increased robustness may have been required for primates to negotiate more difficult unpredictable environments, such as the forest canopy. Direct cortical control of the musculature may have evolved because it allowed primates to avoid their less “dexterous” predators simply by ascending, and robustly negotiating, the precarious branches of tree tops. However, the consequences of this cortical “take-over” might be even more profound.

With motor cortex in direct control of overt movements, the behaviour of a primate is a more direct reflection of cortical state: when you watch a primate move you are directly observing cortical commands. For species that live in social groups, this would allow a uniquely efficient means of communicating the state of cortex between conspecifics, a rather significant advantage for group coordination and a likely prerequisite for human language. This novel role for motor cortex—communication—might have exerted the evolutionary pressure to give cortex increasing control over basic movements, ultimately obscuring its primordial, and fundamental, role in robust control.

### Some preliminary conclusions

Clearly our results are insufficient to draw any final conclusion, but that is not our main goal. We present these experiments to support and motivate our attempt to distil a long history of research, and ultimately suggest a new approach to investigating the role of motor cortex. This approach most directly applies to studies of non-primate mammals. There is now a host of techniques to monitor and manipulate cortical activity during behaviour in these species, but we propose that we should be monitoring and manipulating activity during behaviours that actually require motor cortex.

This synthesis also has implications for engineers and clinicians. We suggest that acknowledging a primary role for motor cortex in robust control, a problem still daunting to robotics engineers, can guide the development of new approaches for building intelligent machines, as well as new strategies to assess and treat patients with motor cortical damage. We concede that our results are still naïve, but propose that the implications are worthy of further consideration.

## Methods

All experiments were approved by the Champalimaud Foundation Bioethics Committee and the Portuguese National Authority for Animal Health, Direcção-Geral de Alimentação e Veterinária (DGAV).

### Permanent lesions

Ibotenic acid was injected bilaterally in 11 Long-Evans rats (ages from 83 to 141 days; 9 females, 2 males), at 3 injection sites with 2 depths per site (−1.5 mm and −0.75 mm from the surface of the brain). At each depth we injected a total amount of 82.8 nL using a microinjector (Drummond Nanoject II, 9.2 nL per injection, 9 injections per depth). The coordinates for each site, in mm with respect to Bregma, were: +1.0 AP / 2.0 ML; +1.0 AP / 4.0 ML; +3.0 AP / 2.0 ML, following the protocol reported by Kawai et al. for targeting forelimb motor cortex (Kawai et al., 2015). Five other animals were used as sham controls (age-matched controls; 3 females, 2 males), subject to the same intervention, but where ibotenic acid was replaced with physiological saline. Six additional animals were used as wildtype, no-surgery, controls (age-matched controls; 6 females).

For the frontal cortex aspiration lesions, the margins of the craniotomy were extended to cover from -2.0 to + 5.0 mm AP relative to Bregma and laterally from 0.5 mm up to the temporal ridge of the skull. After removal of the skull, the exposed dura was cut and removed, and the underlying tissue aspirated to a depth of 2 to 3 mm with a fine pipette (Whishaw, 2000). For the frontoparietal cortical lesions, the craniotomy extended from -6.0 to + 4.0 mm AP relative to Bregma and laterally from 0.5 mm up to the temporal ridge. Two of these animals underwent aspiration lesions as described above. In the remaining animal, the lesion was induced by pial stripping in order to further restrict the damage to cortical areas. After removal of the dura, the underlying pia, arachnoid and vasculature were wiped with a sterile cotton swab until no vasculature was visible (Farr and Whishaw, 2002).

### Recovery period

After the surgeries, animals were given a minimum of one week (up to two weeks) recovery period in isolation. After this period, animals were handled every day for a week, after which they were paired again with their age-matched control to allow for social interaction during the remainder of the recovery period. In total, all animals were allowed at least one full month of recovery before they were first exposed to the behaviour assay.

The three largest frontoparietal lesioned animals were originally prepared for a study of behaviour in a dynamic visual foraging task, which they were exposed to for one month in addition to the recovery period described above. This task did not, however, require any challenging motor behaviours besides locomotion over a completely flat surface. This period was also used to monitor the overall health condition of the animals and to facilitate sensorimotor recovery as much as possible. The animal with the largest lesion (Extended Lesion F) was prevented from completing the behaviour protocol due to deteriorating health conditions following the first two days of testing.

### Histology

All animals were perfused intracardially with 4% paraformaldehyde in phosphate buffer saline (PBS) and brains were post-fixed for at least 24 h in the same fixative. Serial coronal sections (100 µm) were Nissl-stained and imaged for identification of lesion boundaries. In two of the largest frontoparietal lesions (Extended Lesions D and E), serial sections were taken sagittally.

In order to reconstruct lesion volumes, the images of coronal sections were aligned and the outlines of both brain and lesions were manually traced in Fiji (Schindelin et al., 2012) and stored as two-dimensional regions of interest. Lesion volumes were calculated by summing the area of each region of interest multiplied by the thickness of each slice. The stored regions were also used to reconstruct a 3D polygon mesh for visualization of lesion boundaries.

### Electrocorticography

Recording of electrophysiological signals from the intact rodent cortex was performed using two high-density 64-channel micro-electrocorticography (micro-ECoG) grids using the method reported by Dimitriadis et al. for freely moving animals (Dimitriadis et al., 2014). The particular grids used in these experiments were fabricated at the International Iberian Nanotechnology Laboratory by depositing microelectrode gold contacts through a custom designed layout mask on a flexible thin-film polyimide substrate (Figure 11A). The soft connectors at the end of each grid are inserted during the implantation surgery into a custom made breakout board in the recording chamber, which exposes groups of 32-channels via Omnetics connectors to the recording amplifier (see data acquisition section).

The microelectrode grids were implanted epidurally into the right hemisphere of three male Long-Evans rats at almost two years of age. The margins of the craniotomy for implantation extended from -3.3 to + 5.0 mm AP relative to Bregma and laterally from 1.5 to 4.0 mm. The anterior grid was first placed carefully on top of the brain, and then slowly inserted below the anterior and medial margins of the craniotomy until the first rows of electrodes were fully covered. The second grid was placed posterior to the first one and inserted below the medial margin of the craniotomy, taking care that the first rows of electrodes were kept equidistant from the last row of electrodes in the anterior grid. Two zirconium hooks were inserted in the anterior and posterior margins of the craniotomy and fixed to the recording chamber in order to hold it firmly in place relative to the skull. With the aid of a micromanipulator and video feedback system, the coordinates of different electrodes in each quadrant of both grids were measured relative to Bregma, and later used to reconstruct the precise placement of all grid electrodes in the brain. At the end of the surgery, a titanium screw was inserted posteriorly to the craniotomy in contact with the brain in order to be used as reference for the recording system. The stability of the implant depends critically on the absence of movement in the bony plates of the skull during development, which can compromise the mechanical fixation of the recording chamber to the head (Dimitriadis et al., 2014). For this reason, it is recommended that rats undergoing this procedure should be older than 7 months (Dimitriadis et al., 2014).

### Behaviour assay

During each session the animal was placed inside a behaviour box for 30 min, where it could collect water rewards by shuttling back and forth between two nose pokes (Island Motion Corporation, USA). To do this, animals had to cross a 48 cm obstacle course composed of eight 2 cm aluminium steps spaced by 4 cm (Figure 2A). The structure of the assay and each step in the obstacle course was built out of aluminium structural framing (Bosch Rexroth, DE, 20 mm series). The walls of the arena were fabricated with a laser-cutter from 5 mm thick opaque black acrylic and fixed to the structural framing. A transparent acrylic window partition was positioned in front of the obstacle course in order to provide a clear view of the animal. All experiments were run in the dark by having the behavioural apparatus enclosed in a light tight box.

A motorized brake allowed us to lock or release each step in the obstacle course (Figure 2B). The shaft of each of the obstacles was coupled to an acrylic piece used to control the rotational stability of each step. In order to lock a step in a fixed position, two servo motors are actuated to press against the acrylic piece and hold it in place. Two other acrylic pieces were used as stops to ensure a maximum rotation angle of approximately +/− 100°. Two small nuts were attached to the bottom of each step to work as a counterweight that gives the obstacles a tendency to return to their original flat configuration. In order to ensure that noise from servo motor actuation could not be used as a cue to tell the animal about the state of each step, the motors were always set to press against an acrylic piece, either the piece that keeps the step stabilized, or the acrylic stops. At the beginning of each trial, the motors were run through a randomized sequence of positions in order to mask information about state transitions and also to ensure the steps were reset to their original configuration. Control of the motors was done using a Motoruino board (Artica, PT) along with a custom workflow written in the Bonsai visual programming language (Lopes et al., 2015).

Prior to the micro-ECoG recordings, each step in the obstacle course was outfitted with a micro load cell (CZL616C, Phidgets, CA) secured between the step front holder and the base (Figure 2B). This allowed us to record a varying voltage signal proportional to the load applied by the animal on each step. This load signal was acquired simultaneously on all eight steps and digitized synchronously with the ECoG data acquisition system.

### Data acquisition

The behaviour of the animals was recorded with a high-speed and high-resolution videography system (1280 × 680 @ 120 Hz) using an infrared camera (Flea3, PointGrey, CA), super-bright infrared LED front lights (SMD5050, 850 nm) and a vari-focal lens (Fujinon, JP) positioned in front of the transparent window partition. A top view of the assay was simultaneously recorded with the same system at a lower frame-rate (30 Hz) for monitoring purposes. All video data was encoded with MPEG-4 compression for subsequent o[ine analysis. Behaviour data acquisition for the nose poke beam breaks was done using an Arduino board (Uno, Arduino, USA) and streamed to the computer via USB. All video and sensor data acquisition was recorded in parallel using the same Bonsai workflow used to control the behaviour assay.

For the micro-ECoG recordings, all electrophysiological signals were amplified, digitized and multiplexed using two 64-channel amplifier boards (RHD2164, Intan Technologies, US) connected to the electrode interface board (EIB) on the recording chamber. The amplifier boards were then connected through a dual headstage adapter (C3440, Intan Technologies, US) to the main data acquisition USB interface board (RHD2000-Eval, Intan Technologies, US). In order to facilitate the free movement of the animal in the behaviour box, the single cable connecting the head of the animal to the USB interface board was passed through a slip ring (MMC235, Moflon, CN) and hooked into a nylon string crossing the top of the assay. In this way, movement and rotation of the tethered animal were compensated to avoid unwanted strain and twisting on the cables during the entire recording period.

In order to synchronize the videography and ECoG recording systems, we connected the strobe output of the camera to a digital input in the Intan USB interface board using a GPIO cable (ACC-01-3000, PointGrey, CA). The camera strobe output is electronically coupled to individual frame exposures (i.e. shutter opening and closing events), and can be used for sub-millisecond readout of individual frame acquisition times. The strobe signal was acquired and digitized synchronously with ECoG data acquisition, and used for *post-hoc* reconstruction of precise frame timing. Data acquisition from the USB interface board was recorded using a Bonsai workflow and care was taken that it was always started first and terminated last in order to ensure that no external synchronization events were lost.

### Behaviour protocol

The animals were kept in a state of water deprivation for 20 h prior to each daily session. For every trial, rats were delivered a 20 µL drop of water. At the end of each day, they were given free access to water for 10 min before initiating the next deprivation period. Sessions lasted for six days of the week from Monday to Saturday, with a day of free access to water on Sunday. Before the start of the water deprivation protocol, animals were run on a single habituation session where they were placed in the box for a period of 15 min.

The following sequence of conditions were presented to the animals over the course of a month (see also Figure 2A): day 0, habituation to the box; day 1-4, all the steps were fixed in a stable configuration; day 5, 20 trials of the stable configuration, after which the two centre steps were made unstable (i.e. free to rotate); day 6-10, the centre two steps remained unstable; day 11, 20 trials of the unstable configuration, after which the two centre steps were again fixed in a stable state; day 12, all the steps were fixed in a stable configuration; day 13-16, the state of the centre two steps was randomized on a trial-by-trial basis to be either stable or unstable. Following the end of the random protocol, animals continued to be tested in the assay for a variable number of days (up to one week) in different conditions. At the end of the testing period, all animals were exposed to a final session where all steps were made free to rotate in order to assay locomotion performance under challenging conditions.

For the micro-ECoG recordings, the basic behaviour protocol was adjusted to allow for extra recording time during conditions of interest. First, all session times were doubled for the recordings (e.g. 30 min for the habituation session, and 60 min for all other sessions). Second, the number of days on each condition was also extended to allow extracting more trials from each animal for analysis. Finally, the condition where the centre two steps were reliably unstable was replaced with a condition of rare instability. In this condition, after the animal is exposed to an unstable configuration, the steps are reverted back to being stable for another 20 trials, after which they become again unstable for one trial, and so on.

### Data analysis

All scripts and custom code used for data analysis are available online^1^. The raw video data was first pre-processed using a custom Bonsai workflow in order to extract features of interest (Figure 2C). Tracking of the nose was achieved by background subtraction and connected component labelling of segmented image elements. First we compute the ellipse best-fit to the largest object in the image. We then mark the tip of the nose as the furthermost point, in the segmented shape of the animal, along the major axis of the ellipse. In order to analyse stepping performance, regions of interest were defined around the surface of each step and in the gaps between the steps. Background subtracted activity over these regions was recorded for every frame for subsequent detection and classification of steps and slips.

Analysis routines were run using the NumPy scientific computing package (van der Walt et al., 2011) and the Pandas data analysis library (McKinney, 2010) for the Python programming language. Crossings were automatically extracted from the nose trajectory data by first detecting consecutive time points where the nose was positively identified in the video. In order for these periods to be successfully marked as crossings, the starting position of the nose must be located on the opposite side of the ending position. Inside each crossing, the moment of stepping with the forelimb on the centre steps was extracted by looking at the first peak above a threshold in the first derivative of the activation signal in the corresponding region of interest. False positive classifications due to hindlimb or tail activations were eliminated by enforcing the constraint that the position of the head must be located before the next step. Visual confirmation of the classified timepoints showed that spurious activations were all but eliminated by this procedure as stepping with the hindlimb or tail requires the head to be further ahead in space unless the animal turned around (in which case the trajectory would not be marked as a crossing anyway). The position of the nose at the moment of each step was extracted and found to be normally distributed, so statistical analysis of the step posture in the random condition used an unpaired t-test to check for independence of different measurement groups.

In order to evaluate the dynamics of crossing in the random condition, we first measured for every trial the speed at which the animals were moving on each spatial segment of the assay. To minimize overall trial-by-trial variation in individual animal performance, we used the average speed at which the animal approached the manipulated step as a baseline and subtracted it from the speed at each individual segment. To summarize differences in performance between stable and unstable trials, we then computed the average speed profile for each condition, and then subtracted the average speed profile for unstable trials from the average speed profile for stable trials. Finally, we computed the sum of all these speed differences at every segment in order to obtain the speedup index for each animal, i.e. an index of whether the animal tends to accelerate or decelerate across the assay on stable versus unstable trials.

For the micro-ECoG experiments, evoked potentials were analysed by splitting the raw physiological voltage traces into 750 ms windows, where time zero was aligned to the moment of stepping with the forelimb on one of the obstacles in the course (see below). Each individual time series was low-pass filtered at 50 Hz (4th order Butterworth filter, two-pass) and baselined by subtracting the average of the first 250 ms before event onset in order to compensate for constant voltage shifts between the two grids. Some of the channels in each grid were entirely excluded from the analysis due to potentially damaged surface contacts, as evidenced by wide amplitude, random oscillatory behaviour, which was often matched by the presence of high impedance measurements extracted from the electrode site in vivo. In one of the sessions, the cable connecting the headstage to the interface board was accidentally removed by the animal, and all the trials falling during this period had to be excluded from analysis. Correspondence between individual ECoG samples and video frames was computed by matching the individual hardware frame counter with the sequence of falling edges detected in the shutter strobe signal acquired from the infrared camera.

### Video classification

Classification of paw placement faults (i.e. slips) was performed in semi-automated fashion. First, possible slip timepoints were detected automatically using the peak detection method outlined above. All constraints on head position were relaxed for this analysis in order to exclude the possibility of false negatives. A human classifier then proceeded to manually go through each of the slip candidates and inspect the video around that timepoint in order to assess whether the activation peak was a genuine paw placement fault. Examples of false positives include tail and head activations as well as paw activations that occur while the animal is actively engaged in exploration, rearing, or other activities that are unrelated to crossing the obstacles.

A similar technique was used to detect and classify the event onsets for the analysis of evoked potentials in the micro-ECoG experiments. In this case, a preliminary classification of each video frame into left and right forelimb was achieved by first computing the brightness histogram of each frame, which was used to encode the image as a lower-dimensional vector. The vectors for all step frames were subsequently clustered using K-means and then manually inspected for label correction.

Classification of behaviour responses following first exposure to the unstable condition was done on a frame-by-frame analysis of the high-speed video aligned on first contact with the manipulated step. The frame of first contact was defined as the first frame in which there is noticeable movement of the step caused by animal contact. Three main categories of behaviour were observed to follow the first contact: compensation, investigation and halting. Behaviour sequences were first classified as belonging to one of these categories and their onsets and offsets determined by the following criteria. Compensation behaviour is defined by a rapid and adaptive postural correction to the locomotion pattern in response to the perturbation. Onset of this behaviour is defined by the first frame in which there is visible rapid contraction of the body musculature following first contact. Investigation behaviour consists of periods of targeted interaction with the steps, often involving manipulation of the freely moving obstacle with the forepaws. The onset of this behaviour is defined by the animal orienting its head down to one of the manipulated steps, followed by subsequent interaction. Halting behaviour is characterized by a period in which the animal stops its ongoing motor program, and maintains the same body posture for several seconds, without switching to a new behaviour or orienting specifically to the manipulated steps. This behaviour is distinct from a freezing response, as occasional movements of the head are seen. Onset of this behaviour is defined by the moment where locomotion and other motor activities besides movement of the head come to a stop. A human classifier blind to the lesion condition was given descriptions of each of these three main categories of behaviour and asked to note onsets and offsets of each behaviour throughout the videos. These classifications provide a visual summary of the first response videos; the complete dataset used for this classification is included as supplementary movies.

## Author contributions

Conceived and designed the lesion experiments: G.L., A.R.K., J.J.P.; Performed the lesion experiments: G.L., J.N.; Analysed the lesion data: G.L., A.R.K.; Conceived and designed the ECoG experiments: G.L., G.D., A.R.K., J.J.P.; Performed the ECoG implantation surgeries: G.D., J.N.; Performed the ECoG behaviour experiments: G.L.; Analysed the ECoG data: G.L., G.D., J.A.M., A.R.K.; Wrote the manuscript: G.L., A.R.K.

## Competing interests

The authors declare that the research was conducted in the absence of any commercial or financial relationships that could be construed as a potential conflict of interest.

## Acknowledgements

We thank Lorenza Calcaterra for the extended frontoparietal cortical lesion preparations; João Gaspar of the International Iberian Nanotechnology Laboratory for kindly providing the fabrication process for the micro-ECoG grids; João Frazão, Pedro Lacerda and Tiago Monteiro for invaluable help in extending the behaviour assay for the micro-ECOG recordings and all the members of the Intelligent Systems Lab for constant feedback on the ideas, experiments and manuscript as well as help annotating behaviour videos. The research leading to these results has received funding from the European Union’s Seventh Framework Programme (FP7/2007-2013) under grant agreement no. 600925. G.L. was supported by the PhD Studentship SFRH/BD/51714/2011 from the Foundation for Science and Technology. The Champalimaud Neuroscience Programme is supported by the Champalimaud Foundation.

## Footnotes

1 https://bitbucket.org/kampff-lab/shuttling-analysis

